# Seasonal dynamics and environmental drivers of tissue and mucus microbiomes in the staghorn coral *Acropora pulchra*

**DOI:** 10.1101/2023.09.07.556622

**Authors:** Therese C Miller, Bastian Bentlage

## Abstract

**Background:** Rainfall-induced coastal runoff represents an important environmental impact in near-shore coral reefs that may affect coral-associated bacterial microbiomes. Shifts in microbiome community composition and function can stress corals and ultimately cause mortality and reef declines. Impacts of environmental stress may be site specific and differ between coral microbiome compartments (e.g., tissue *versus* mucus). Coastal runoff and associated water pollution represent a major stressor for near-shore reef-ecosystems in Guam, Micronesia.

**Methods:** *Acropora pulchra* colonies growing on the West Hagåtña reef flat in Guam were sampled over a period of eight months spanning the 2021 wet and dry seasons. To examine bacterial microbiome diversity and composition, samples of *A. pulchra* tissue and mucus were collected during late April, early July, late September, and at the end of December. Samples were collected from populations in two different habitat zones, near the reef crest (farshore) and close to shore (nearshore). Seawater samples were collected during the same time period to evaluate microbiome dynamics of the waters surrounding coral colonies. Tissue, mucus, and seawater microbiomes were determined using 16S DNA metabarcoding using Illumina sequencing. In addition, water samples were collected to determine fecal indicator bacteria (FIB) concentrations, as an indicator of water pollution. Water temperatures were recorded using data loggers and precipitation data obtained from a nearby rain gauge. The correlation structure of environmental parameters (temperature and rainfall), FIB concentrations, and *A. pulchra* microbiome diversity was evaluated using a struictural equation model. Beta diversity analyses were used to investigate spatio-temporal trends of microbiome composition.

**Results:** *A. pulchra* microbiome diversity differed between tissues and mucus, with mucus microbiome diversity being similar to the surrounding seawater. Rainfall and associated fluctuations of FIB concentrations were correlated with changes in tissue and mucus microbiomes, indicating their role as drivers of *A. pulchra* microbiome diversity. *A. pulchra* tiussue microbiome composition remained relatively stable throughout dry and wet seasons and were dominated by Endozoicomonadaceae, coral endosymbionts and putative indicators of coral health. In nearshore *A. pulchra* tissue microbiomes, Simkaniaceae, putative obligate coral endosymbionts, were more abundant than in *A. pulchra* colonies growing near the reef crest (farshore). *A. pulchra* mucus microbiomes were more diverse during the wet season than the dry season, a distinction that was also associated with drastic shifts in microbiome composition. This study highlights the seasonal dynamics of coral microbiomes and demonstrates that microbiome diversity and composition may differ between coral tissues and the surface mucus layer.

## Introduction

Coral reefs are largely restricted to shallow waters that are strongly affected by environmental conditions that may vary across small spatial scales such as sea surface temperatures, water flow, and tidal ranges (Guilcher, 1988). Shallow near-shore coral reefs are also susceptible to anthropogenic impacts such as pollution and nutrient runoff (Hughes, 1994). Pollution by sewage and coastal runoff is a significant source of environmental stress that negatively impacts coral reefs by disrupting coral microbiome communities and their interactions with their coral host (Wooldridge and Done, 2009). Sewage is highly enriched in ^15^N and has a distinct stable isotope composition in comparison to other nitrogen sources and can be detected in, for example, macroalgae as a signature of long-term pollution (Abaya et al., 2018). On Hawaiian reefs, percent coral cover was negatively correlated with nitrogen pollution, including macroalgal-incorparted δ^15^N and elevated concentrations of fecal indicator bacteria (FIB) (Abaya et al., 2018). In Guam, Redding et al. (2013) found a positive correlation between sewage-derived nitrogen and severity of coral disease as well as a correlation between precipitation and the concentration of δ^15^N in near-shore reefs. In addition to being a source of nitrogen and eutrophication, sewage and runoff carry bacteria into near-shore coral reef ecosystems (Sutherland et al., 2011; Haapkylä et al., 2011), potentially disrupting coral-associated bacterial microbiomes (Maher, Epstein and Vega Thurber, 2022). For example, bacterial microbiome diversity was shown to be elevated in corals exposed to sewage and municipal wastewater (Ziegler et al., 2016).

Bacteria are integral parts of the coral holobiont and are involved in nutrient cycling and the mitigation of pathogens (Maher et al., 2022; Bourne et al., 2016). Bacterial microbiomes may differ between compartments of the coral, including the skeleton, tissue, and the surface muco-polysaccharide layer (mucus hereafter) (Glasl et al., 2016; Hernandez-Agreda et al., 2017; Peixoto et al., 2017). As it is in direct contact with the environment, the mucus is a first line of defense for corals against pathogens and plays an important role in disease mitigation through antibiotic activity (Hernandez-Agreda et al., 2017; Ritchie, 2006). Depletion of mucus bacteria has been linked to increases in *Vibrio* spp. and other pathogenic bacteria (Glasl et al., 2016). Disturbing the mucus microbiome may thus open the way for pathogens to enter coral tissues and promote disease that may potentially lead to coral mortality (Glasl et al., 2017). The mucus is a dynamic environment and the micro-associates within it are directly exposed to biotic and abiotic variations in the environment, suggesting that the mucus community may be highly variable over time (Hernandez-Agreda et al. 2017). By contrast, the bacterial microbiome of the tissue layer has been shown to be more stable (Marchioro et al., 2020).

Coral bacterial microbiome composition and diversity may be site specific, differing between habitats (Camp et al., 2020). The observation of such site-associated differences in coral microbiomes has led to the classification of some corals as conformers whose microbiomes adapt to the surrounding environment (Voolstra and Ziegler, 2020). Microbiome regulators, by contrast, maintain a consistent microbiome regardless of differences in external environment across sites (Voolstra and Ziegler, 2020). *Porites cylindrica*, a microbiome regulator, exposed to fish farm effluent showed increased abundances of *Vibrio* spp. in its microbiome after five days of exposure, but microbiomes returned to their original states after roughly three weeks of exposure to effluent (Garren et al., 2009). Microbiome conformers such as *Acropora* spp. tend to absorb bacteria from the surrounding environment and incorporate them into their microbiomes (Greer et al., 2009; DeVantier et al., 2006; Ziegler et al., 2019). Staghorn *Acropora* spp., in particular *Acropora pulchra*, dominate Guam’s reef flats, forming thickets whose extent has been reduced over the last decade by a combination of coral bleaching and extreme low tide exposure (Raymundo et al., 2017, 2019, 2022).

Thus far, little is known about the bacterial microbiomes of Guam’s *Acropora pulchra* populations and the impact seasonal runoff may have on microbiome composition. We employed DNA metabarcoding to investigate the seasonal dynamics of *Acropora pulchra* bacterial microbiomes in Guam’s West Hagåtña Bay. To identify possible differences in microbiome composition across coral compartments, microbiomes were characterized separately for both tissue and mucus. Specifically, the objectives of this study were to (1) characterize bacterial microbiome communities of coral tissue and mucus of *A. pulchra* from near-shore and far-shore zones of West Hagåtña Bay and (2) to elucidate the impacts of seasonal change on coral bacterial microbiomes. We found that the microbiomes of *A. pulchra* differed across compartments, with mucus microbiomes being more diverse than tissue microbiomes. The mucus microbiome behaved more like a conformer, reflecting seasonal environmental changes, compared to tissue microbiomes that remained relatively stable throughout the year, similar to a microbiome regulator. Nonetheless, impacts of seasonally varying bacterial pollution on both mucus and tissue microbiome diversity were significant.

## Materials & Methods

### Field Site

West Hagåtña Bay, Guam (Fig. 1) can be divided into two zones: outer (closer to the reef crest) and inner (closer to the shore). These zones correspond to different environmental regimes (Fifer et al., 2021) that saw drastically different mortality rates during past coral bleaching and extreme low tide events (Raymundo et al., 2017). In each zone, three representative sites of *Acropora pulchra* stands were selected for sampling. At each site, three coral colonies were tagged to ensure repeated sampling of the same coral colonies (Fig. 2). All coral collections for this study were made under a special license for the collection of coral issued by Guam’s Department of Agriculture to the University of Guam Marine Laboratory.

**Figure 1.**
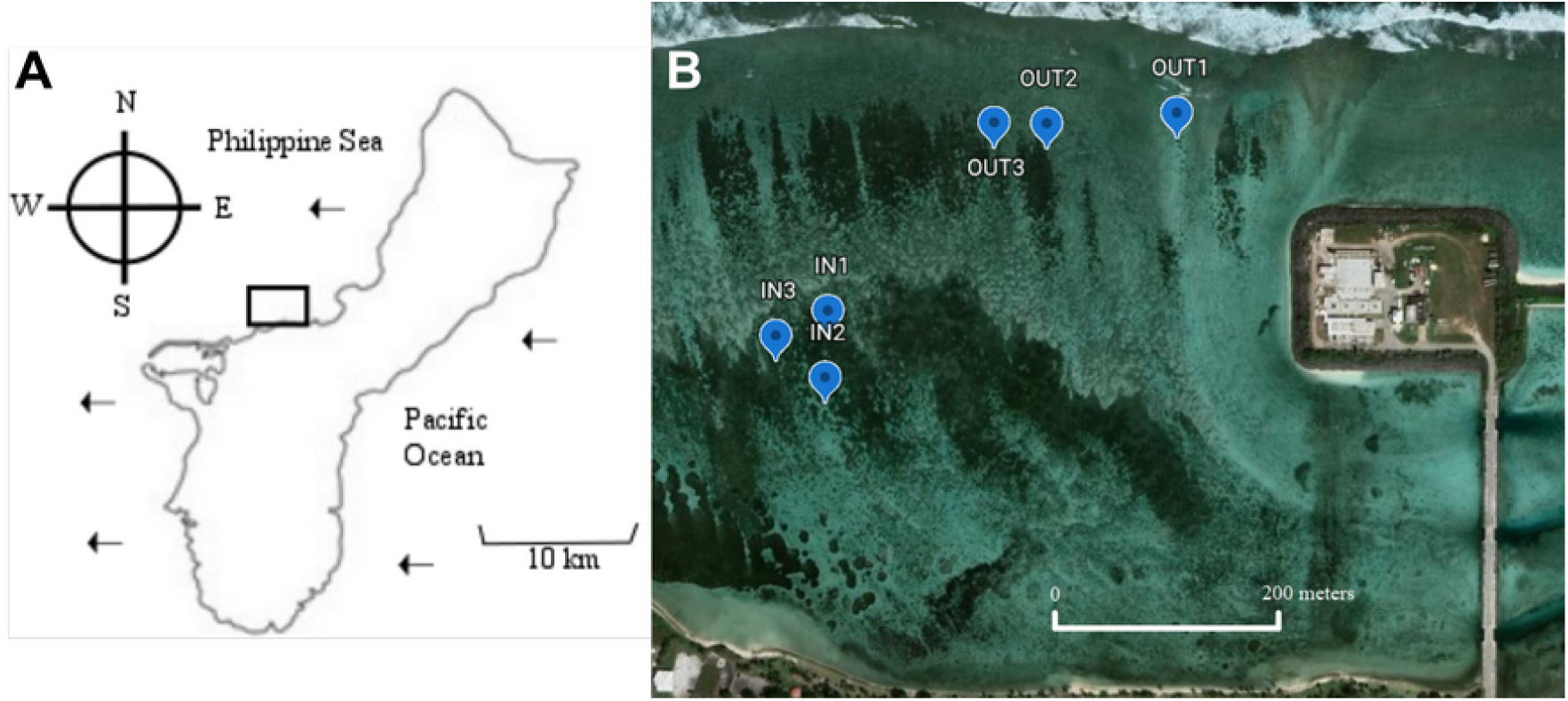
Sampling sites targeted during this study. West Hagatna Bay’s location in Guam is highlighted by the black rectangle (A). Sites in the inner(IN1-3)and outer(OUT1-3) zones where *Acroporapulchra* colonieswere tagged for repeated sampling are indicated by blue markers. Latitude and Longitude for sampling locations are provided in Supplemental Table 1.

**Figure 2.**
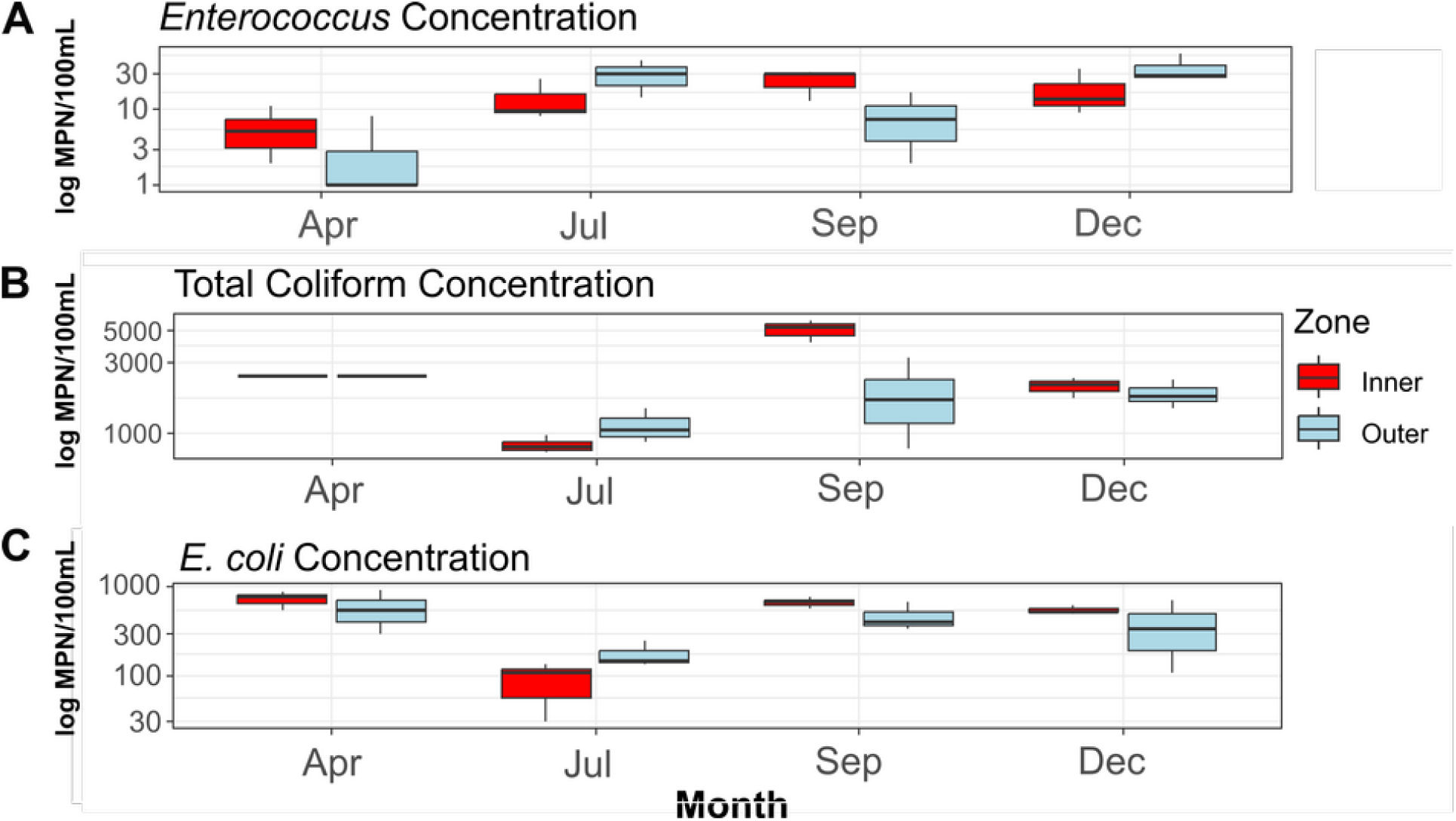
Concentrations of fecal indicator bacteria (FIB) contained in water samples collected during the course of the study. Concentrations of *Enterococcus* (A), total coliform bacteria (B; total coliform concentrations for the month of May provided as > 2419.6 MPN/100 mL), and concentrations of *E. coli* (C) are provided as log10-scaled most probable numbers (MPN).

### Environmental data

Six HOBO TidBit temperature loggers (Onset Corp., Bourne, MA) were deployed at 1m depth, one at each sampling site, to record seawater temperatures in 5 minute intervals from April to December 2021, the duration of this study. Daily precipitation data were collected by the National Oceanic and Atmospheric Administration (NOAA) at the Guam International Airport and retrieved from the NOAA website (NOAA, 2022). During each of the four microbiome specimens collection events (see below), seawater samples were collected from each site using sterile 50 ml polypropylene centrifuge tubes. Fecal indicator bacteria (FIB) concentrations (*Escherichia coli*, *Enterococcus*, and total coliform) were quantified by the Water and Environmental Research Institute (WERI) at the University of Guam. Separate 50 ml water samples were collected for determination of nitrate/nitrite (NO ^-^/NO ^-^) and ortho-phosphate (PO ^3-^) concentrations by WERI.

Temperatures were averaged by day within each zone (inner and outer), and daily averages were used in statistical analyses. Concentrations of FIB were averaged by month for each zone (inner and outer). Daily precipitation obtained from NOAA (2022) was averaged by month for further analysis. Normality of each environmental dataset was estimated using a Shapiro-Wilk test. Two-way ANOVAs incorporating zones and months as factors were used to test for significant differences in FIB concentrations between inner and outer zones as well as between months. A Kruskal-Wallis Chi^2^ test was used to test for significant differences in rainfall per month. Seawater temperature differences between inner and outer zones as well as different months were tested for significance using Kruskal-Wallis Chi^2^ tests on temperature.

### Microbiome sample collection

Samples were collected twice during Guam’s dry season, at the beginning of May and beginning of July 2021, and twice in Guam’s wet season, at the end of September and the end of December 2021. Coral nubbins (4.5 – 5 cm) were sampled from each of the nine tagged *A*. *pulchra* colonies in the outer zone and the nine tagged colonies in the inner zone. Sampling four times throughout the year yielded a total of 72 coral tissue and mucus samples, respectively.

The terminal end of each cut coral fragment was removed since new tissue layers at the growing tip may not be fully colonized by the coral microbiome community. The mucus was sampled by exposing the cut coral nubbin to air and swabbing mucus from tissues using sterilized cotton swabs (Lampert et al., 2008). Samples were frozen in the field using liquid nitrogen and stored at -80°C prior to DNA extraction and PCR.

To characterize the bacterial microbiome of the seawater in West Hagåtña Bay, 3L of seawater were collected during each collection event from each site. Seawater was stored in wide mouth plastic jars that had been sterilized for at least 30 minutes by soaking in 10% bleach solution prior to sample collection. Seawater was transported to the University of Guam Marine Laboratory and filtered using a 1.2 μm nylon filter (Sigma-Aldrich, St Louis, MO) to collect bacteria for DNA extraction. Nylon filters were placed in Whirl-Pak sample bags (Filtration Group, Oakbrook, IL) and snap frozen in liquid nitrogen prior to storage at -80°C.

### DNA extraction and metabarcoding

DNA was extracted from coral tissue, mucus, and seawater samples using the DNEasy Powersoil kit (Qiagen, Hildenheim, Germany) following the manufacturer’s protocol. DNA concentrations were quantified using Qubit fluorometric quantification (Thermo Fisher Scientific, Waltham, MA). Tissue DNA extracts were diluted to a concentration of 10 ng/μl; mucus and seawater extracts yielded around 1 ng/μl of DNA and were not further diluted. The V4 hypervariable region of 16S ribosomal DNA was amplified from each sample using universal bacterial primers primers 515F (Walters et al., 2016) and modified 806R (Apprill et al., 2015). The 30-μl PCR reactions included 3 μl of template DNA, PCR-grade water, 1x ExTaq buffer (Takara Bio, San Jose, CA), 2.5 mM dNTPs, 10 μM forward and 10 μM reverse primers, and 0.75 U ExTaq DNA Polymerase (Takara Bio, San Jose, CA). The thermocycler protocol comprised 30 cycles of initial denaturation at 95°C for 40 seconds, annealing at 58°C for two minutes, and extension at 72°C for one minute; final elongation was performed at 72°C for 5 minutes. A negative control using PCR-grade water instead of DNA as a template was included in every PCR run. All PCR products were checked on a 1% agarose gel stained with GelRed (Sigma-Aldrich, St Louis, MO). PCR products that yielded bright fluorescent bands were considered successful. A subset of mucus samples that produced low yields from the initial PCR were re-amplified. PCR products were purified using the GeneJet PCR Purification Kit (Thermo Fisher Scientific, Waltham, MA) following the manufacturer’s protocol and quantified using Qubit fluorometric quantification (Thermo Fisher Scientific, Waltham, MA).

Purified PCR products were barcoded using indexes and MiSeq adapters synthesized by Macrogen (Seoul, Republic of Korea). Each indexing PCR reaction contained 2 μl of PCR product, 1 mM MiSeq adapter, PCR-grade water, 1x ExTaq buffer, 2.5 mM dNTPs, and 0.5 U ExTaq DNA polymerase per 20 μl reaction. Adapters were ligated to PCR products using five PCR cycles, including denaturation at 95°C for 40 seconds, annealing at 59°C for two minutes, and an extension at 72°C for one minute; a final elongation step at 72°C for seven minutes followed the five cycles. PCR products were pooled, resulting in two sequencing libraries that were purified using a GeneJet PCR Purification Kit (Thermo Fisher Scientific, Waltham, MA), followed by resuspension of each library in 40 μl of elution buffer; each library pool contained a mixture of sampling timepoints to mitigate potential batch effects. 10 μl of each sequencing library were run through a 2% TBE agarose gel and the target band representing the sequencing library was excised using a sterilized scalpel. 200 μl of nuclease-free water were added to the excised gel band and incubated overnight at 4°C. DNA in the resulting solution was purified using the GeneJet PCR Purification Kit (Thermo Fisher Scientific, Waltham, MA) and resuspended again in 20 μl of elution buffer. The size distribution of the purified sequencing library was verified using an Agilent 4150 TapeStation (Agilent, Santa Clara, CA) with a D1000 ScreenTape assay. Libraries were sequenced at CD Genomics (Shirley, NY) using Illumina MiSeq (Illumina, San Diego, CA) sequencing yielding 300 bp paired-end reads.

### Sequence data quality control and taxonomic assignment

The R package DADA2 (Callahan et al., 2016) was used to remove primer sequences, truncate reads, calculate error rates, de-duplicate reads and infer amplicon sequence variants ASVs) after merging of paired reads and removal of chimeras. Non-bimeric ASVs were assigned a taxonomy from the Silva v138 dataset (Glöckner et al., 2017) using a naive Bayesian classifier (Wang et al., 2007) with a minimum bootstrap confidence of 50. The phyloseq package (McMurdie and Holmes, 2013) was used to remove ASVs whose taxonomy matched “Mitochondria,” “Chloroplast”, or “Eukaryota”. MCMC.OTU (Matz, 2016) was used to remove ASVs representing < 0.1% of count data and to identify putative outlier samples (those that had total counts falling below a z-score cutoff of -2.5). Samples with fewer than 1,000 reads (n = 2) were discarded and not used in analyses.

### Spatiotemporal variation of alpha diversity

Observed ASV diversity and Shannon diversity were calculated for each microbiome compartment (tissue, mucus, seawater) using the phyloseq package (McMurdie and Holmes, 2013). Evenness was calculated for each compartment by dividing the Shannon diversity index by the natural logarithm of observed diversity. Faith’s Phylogenetic Diversity (PD; Armstrong et al., 2021) was calculated using the picante package in R (Kembel et al., 2010). Normality of diversity metrics for each compartment was tested using a Shapiro-Wilk test. For normally distributed diversity metrics, a two-way analysis of variance (ANOVA) was used to test for differences in alpha diversity across compartments, between months, and between inner and outer zones. Non-parametric analysis of variance (Kruskal-Wallis Chi^2^ test) was used to compare alpha diversity when normality was rejected. A post-hoc Dunn’s Test was then used for multiple comparisons.

### Structural equation modeling

To analyze the effects of environmental factors on microbiome diversity, structural equation models (SEMs) based on d-separation tests (Shipley, 2009) were created using the piecewise SEM package (Lefcheck, 2016). In particular, the relationships between precipitation, temperature, FIB concentrations, and microbiome diversity were evaluated. Linear models (LM) with Shannon diversity as the response variable and environmental variables, including precipitation, temperature, and FIB concentrations as the predictors formed the foundation of SEMs. The relations between FIB concentrations with precipitation and temperature were modeled using linear mixed effects models (LME). Precipitation and temperature were included as fixed factors and zone (inner and outer) was included as a random factor. Tissue and mucus microbiomes maintained a ratio of sample number to predictor variables (d) greater than 5, which is a widely used guideline for inclusion of predictor and response variables in structural equation models (Lefcheck, 2016). Given the low sample size for seawater microbiomes compared to predictor variables (d = 3.83), seawater was removed from consideration for final SEMs.

SEMs linked two exogenous variables (temperature and precipitation), three endogenous variables (FIB concentrations of *Enterococcus*, *E.coli*, and total coliform) and one response variable (Shannon diversity). SEMs tested for effects of (1) precipitation on concentrations of *Enterococcus,* total coliform, and *E. coli*, (2) temperature on concentrations of *Enterococcus*, total coliform, and *E. coli*, and (3) concentration of each FIB on microbiome diversity and the remaining two concentrations of FIBs (e.g., concentration of total coliform on mucus microbiome diversity and concentrations of *E. coli* and *Enterococcus*). Testing for collinearity among FIB concentrations required inclusion of correlated error structures between each FIB. Separate SEMs that included the aforementioned tests were constructed for each compartment of the bacterial microbiome to further explore potential drivers of microbiome diversity.

### Beta diversity and differentially abundant taxa

PERMANOVA tests using the Adonis2 function in the vegan package (Oksanen et al., 2020) were run with 1,000 permutations to calculate beta diversity between microbiome compartments, sampling months, and inner and outer zones based on weighted UniFrac distances. These were visualized using a Principal Coordinate Analysis (PCoA). Weighted UniFrac distances were used in these analyses because it integrates abundance information for ASVs in distance calculations (Lozupone and Knight, 2005). Post hoc pairwise PERMANOVAs were used to identify possible differences between individual months or compartments.

The R package ANCOMBC (Lin and Peddada, 2020) was used to identify bacterial taxa that were differentially abundant between compartments. Considering that the mucus represents the interface in direct contact with both seawater and coral tissue, comparisons were made between mucus and seawater microbiomes as well as between mucus and tissue microbiomes. Taxa identified differentially abundant with a false discovery rate < 0.05 were considered statistically significant.

## Results

### Environmental Data

Water temperatures ranged from roughly 27°C to 35°C (Supplemental Fig. 1). On average, the inner zone was warmer than the outer zone (Supplemental Fig. 1). During the hottest part of the year, from June to August, there was on average a roughly 0.5°C difference between inner and outer zones. However, the temperature difference observed between the two zones was not significant (X^2^ = 3.181, df =1, p = 0.075). Water temperature did differ significantly when comparing across months throughout the year (X^2^ = 275.51, df = 8, p < 0.001). The amount of precipitation significantly increased by our third sampling timepoint in September (X^2^ = 12.39, df = 3, p = 0.006; Supplemental Fig. 2), as is typical of Guam’s wet season.

Average concentrations for each FIB (Fig. 2) did not differ significantly between zones (*Enterococcus*: F = 0.216, df = 1, p = 0.667; total coliform: F = 0.47, df = 1, p = 0.531; *E. coli*: F = 0.258, df = 1, p = 0.638), between sampling months (*Enterococcus:* F = 3.717, df = 1, p = 0.126; total coliform: F = 0.01, df = 1, p = 0.927; *E. coli*: F = 0.068, df = 1, p = 0.807), or when taking the interactions of zones and months into account (*Enterococcus:* F = 0.300, df = 1, p = 0.613; total coliform: F = 0.11, df = 1, p = 0.756; *E. coli*: F = 0.025, df = 1, p = 0.882). No significant amounts of nitrite/nitrate or orthophosphate above a threshold of 0.01 mg/L were detected.

### Microbiome diversity

16S metabarcoding data were deposited in NCBI GenBank’s Sequence Read Archive (SRA; https://www.ncbi.nlm.nih.gov/sra) (Supplemental Table 1). After removal of low abundance ASVs (present in less than 0.1% of samples) and outliers from the 16,949 ASVs initially identified, 321 ASVs were retained for downstream analysis. Observed and phylogenetic ASV diversity was highest in seawater samples, with both tissue and mucus microbiomes being less diverse (Figs. 3A & B). By contrast, Shannon diversity and evenness of mucus microbiomes was similar to seawater sample diversity and evenness, higher than for tissue samples (Figs. 3C & D). Shannon diversity did not show significant differences between inner and outer zones but marginally significant variation across months (Supplemental Table 2). Most interaction terms (month * compartment, zone * compartment, and month * zone * compartment), however, were significant in explaining variation in Shannon diversity (Supplemental Table 2). Zone or month alone did not explain the observed variation but all interaction terms including compartment were significant in explaining variation in evenness (Supplemental Table 3). While variation of phylogenetic diversity of microbiomes was not explained by differences between zones alone, phylogenetic diversity was affected by month, compartment, and all interaction terms of zone, month, and compartment (Supplemental Table 4). Interestingly, Shannon diversity was neither explained by zone or month for tissue microbiomes (Supplemental Table 2) but the interactions of zone and month explained tissue microbiome phylogenetic diversity (Supplemental Table 4) and microbiome evenness (Supplemental Table 3) with month alone also explaining microbiome evenness (Supplemental Table 3; Supplemental Fig. 3). Zone, month, or their interaction did explain the variation of mucus microbiome diversity and evenness, by contrast (Supplemental Tables 2 - 4; Supplemental Fig. 4); month was the only variable that explained seawater microbiome variation, having a significant association with phylogenetic diversity (Supplemental Table 4; Supplemental Fig. 5).

**Figure 3.**
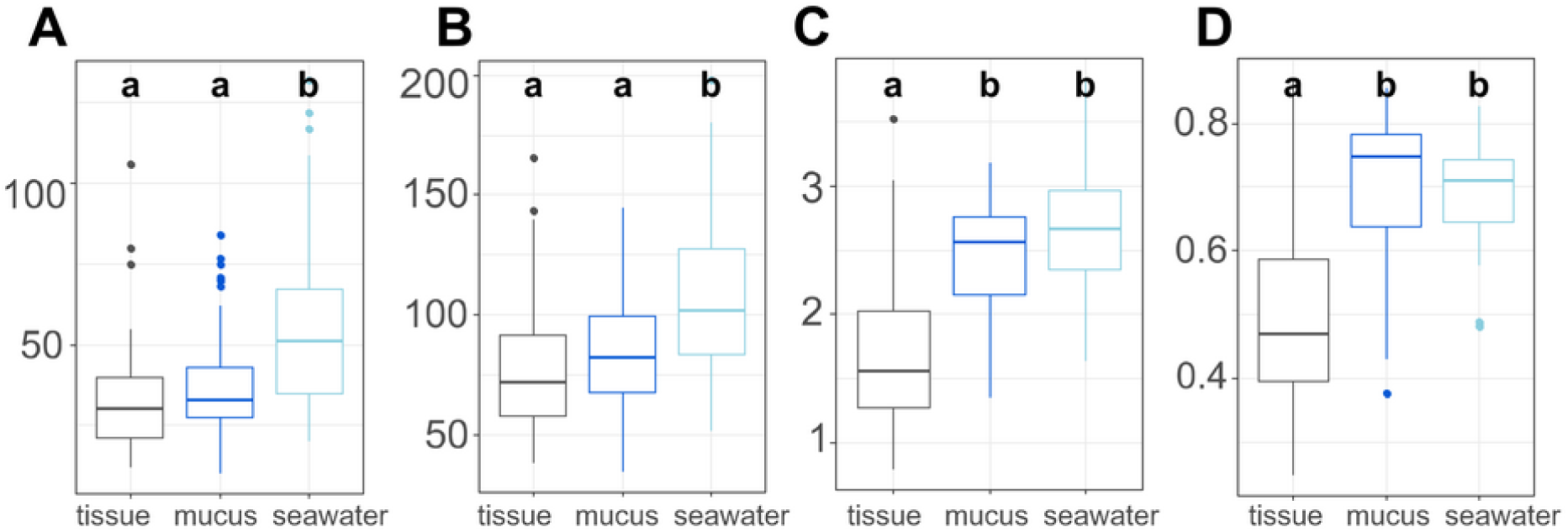
Microbiome diversity and evenness of coral tissues, mucus, and seawater. Boxplots depict minimum, first and third quartile, maximum, and median for observed diversity (A), phylogenetic diversity (B), Shannon diversity (C), and evenness (D). Statistical groups (a and b) were identified using post-hoc Dunn’s tests.

### Environmental predictors of microbiome diversity

Linear mixed effects model analyses for SEMs found the following statistically significant (p < 0.05) linkages: (1) an effect of temperature on FIB concentrations (total coliform, *E.coli*, and *Enterococcus*), (2) an effect of precipitation on FIB concentrations, (3) *E. coli* concentration is subject to common sources of variation with total coliform concentration and *Enterococcus* concentration, (4) an effect of total coliform concentration on tissue microbiome diversity; and (5) an effect of *E. coli* concentration on mucus microbiome diversity (Fig. 4). Temperature and FIB concentrations were negatively correlated (red arrows; Fig. 4) while all other correlations were positive. Total coliform concentration was a significant predictor of tissue microbiome diversity (p = 0.044, R^2^ = 0.09) while *E. coli* concentrations were significant predictors for mucus microbiome diversity (p = 0.019, R^2^ = 0.09). Both of these predictors were weakly associated with their responses, as indicated by the low R^2^ values. Though inclusion of a correlated error structure between precipitation and mucus microbial diversity was not found to be statistically significant (p = 0.089), and precipitation is not hypothesized to directly influence microbial diversity, inclusion of a correlated error structure greatly improved the model’s goodness-of-fit (p = 0.032 to p = 0.637).

**Figure 4.**
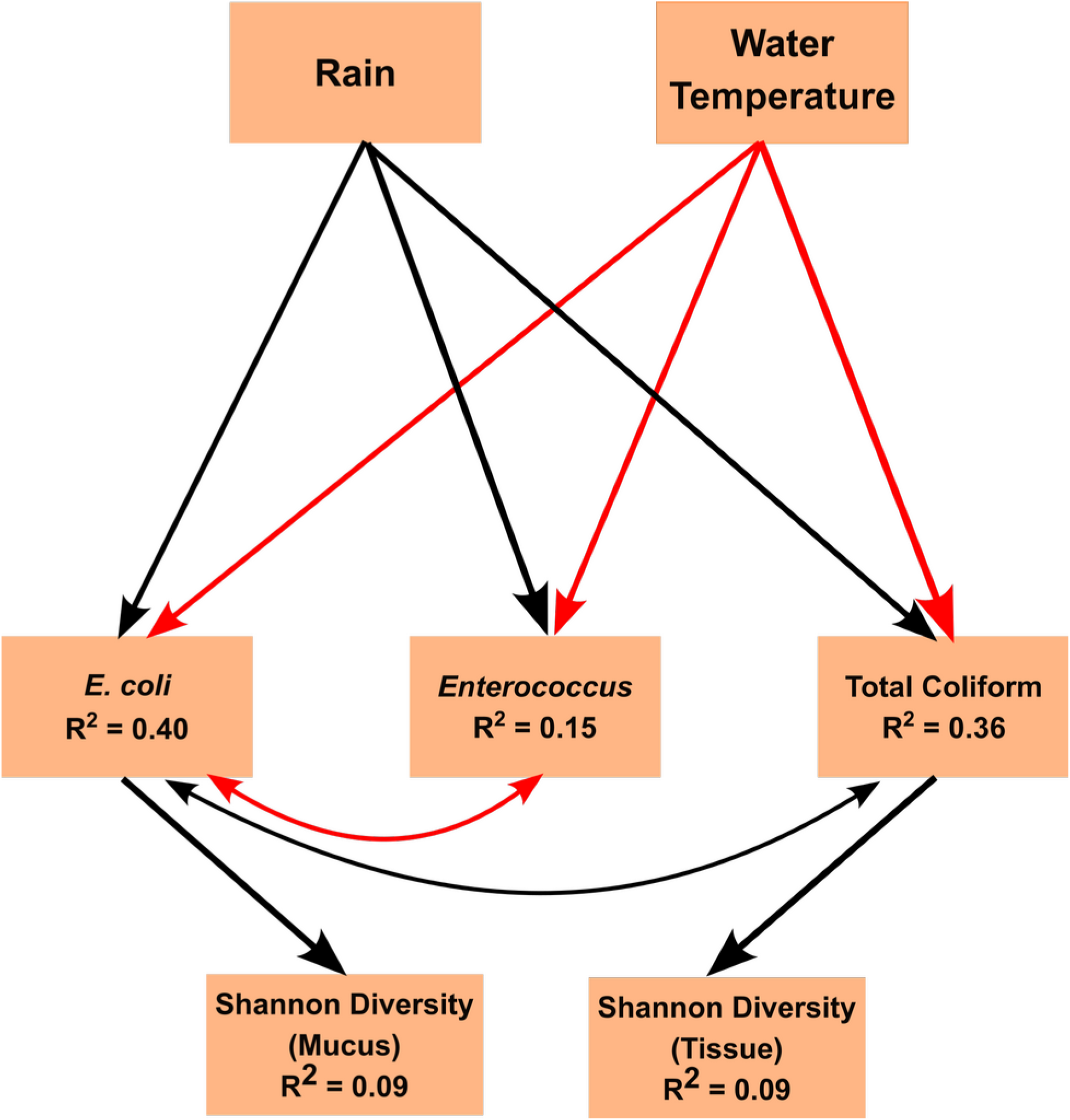
Structural Equation Model (SEM) explaining the relationships between environmental parameters, fecal indicator bacertia (FIB) concentrations, and microbiome diversity. The SEM shows the impacts of exogenous variables (Rain and Temperature) on endogenous variables (concentrations of *E. coli*, *Enterococcus*, and total coliform concentrations) and response variables (Shannon diversity of tissue and mucus). R2 values are given for each endogenous and response variable. Arrows indicate statistically significant (p < 0.05) linkages. Red lines indicate a negative correlation. Black lines indicate a positive correlation.

Arrows shown in the SEMs were supported by Fisher’s C statistics (tissue: C = 5.837, df = 4, p = 0.212; mucus: C = 0.901, df = 2, P = 0.637). The fit of the model to the data could not be rejected by a global goodness-of-fit test (C = 1.972, df = 4, p = 0.741), indicating that no important pathways between variables in the SEM were excluded.

### Microbiome community composition

ASVs associated with the phylum Proteobacteria were dominant overall (Supplemental Table 7), followed by Verrucomicrobiota and Bacteroidota. Among Proteobacteria, the family Endozoicomonadaceae was most abundant, especially in tissue microbiomes (Supplemental Table 8; Fig. 5). In tissue microbiomes, Endozoicomonadaceae dominated all months and zones (Fig. 5A). While Endozoicomonadaceae were dominant in the mucus in April and July during the dry season, their abundance and prevalence dropped sharply in September and December during the wet season (Fig. 5B). Simkaniaceae, Moraxellaceae and Comamonadaceae were the other most abundant taxa in tissue and mucus microbiomes (Fig. 5A & 5B). While Simkaniaceae were more abundant in tissues compared to the mucus, Comamonadaceae and Moraxellaceae were more abundant and prevalent in the mucus compared to tissues (Supplemental Table 8, Fig. 5A and 5B). Endozoicomonadaceae were also found in abundance in seawater samples, in addition to Cyanobiaceae, Rhodobacteraceae, and Chitinophagaceae (Supplemental Table 8, Fig. 5C).

**Figure 5.**
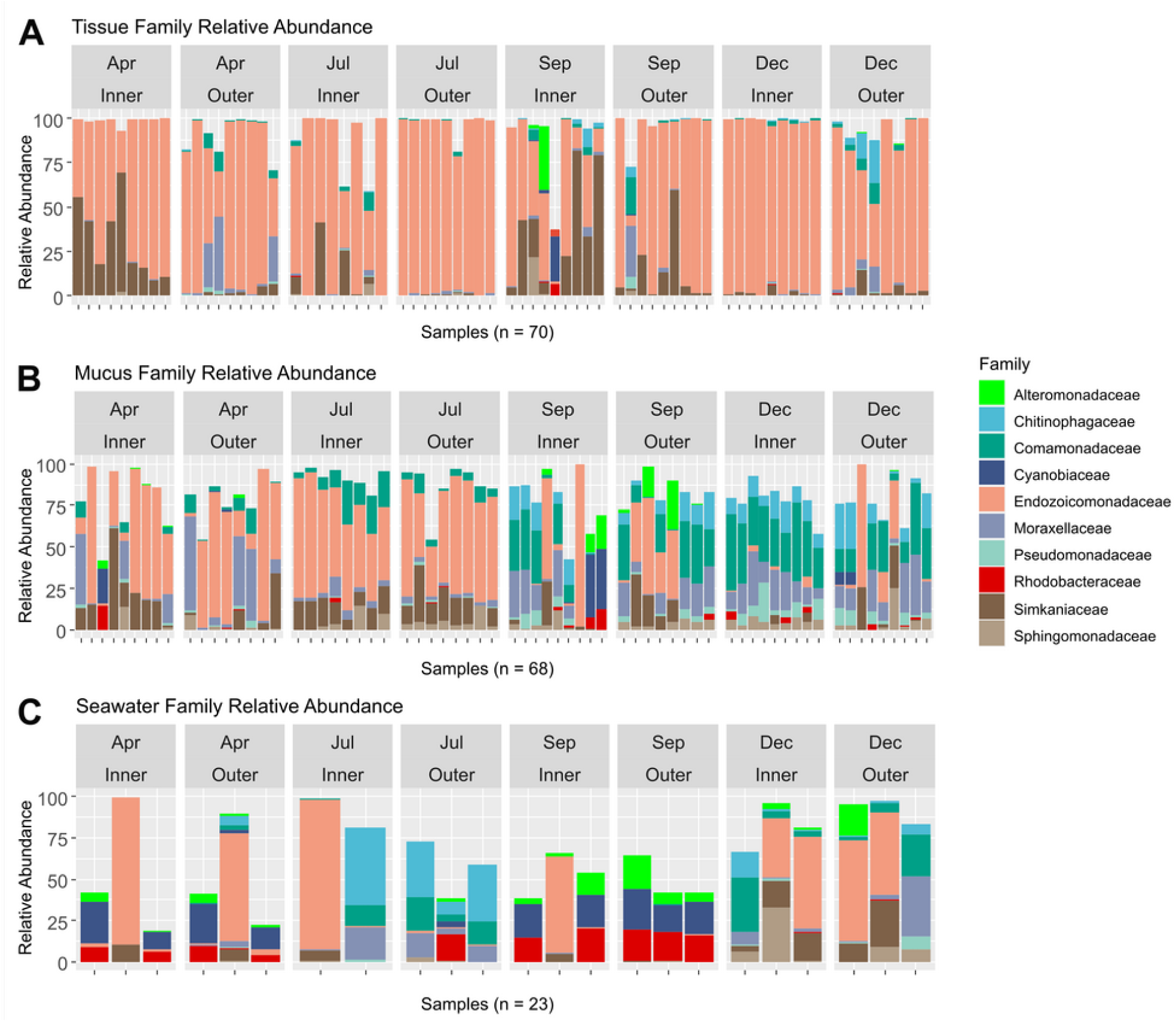
Relative abundances of bacterial taxa identified in coral tissue, mucus, and seawater microbiomes. The ten most abundant bacterial families found in the coral tissue (A), coral mucus (B), and seawater (C) are shown. Samples were separated by months and zones in which they were collected. April and July fell in Guam’s dry season, and September and December fell in Guam’s wet season.

Two PCoA plots based on weighted UniFrac distances were constructed to show differences between compartments, zones, and months (Fig. 6). These plots show the significant differences in microbiome community composition between the different compartments. Additionally, significant differences were seen between months, but not between inner and outer zones (Supplemental Table 5). Some interactions were significant (zone * month, compartment * month), but not all (compartment * zone, compartment * zone * month). Within tissue, there was a significant difference in beta diversity between inner and outer zones, as well as an interaction effect of zones and months, but not between months alone. For the mucus, there was no significant difference between inner and outer zones, or between the interaction of zones and months, but months differed significantly (Supplemental Table 5), as seen with the clustering of mucus samples in the wet season (Fig. 6). In the seawater, no statistically significant difference was seen by month, zone, or the interaction effects (Supplemental Table 5).

**Figure 6.**
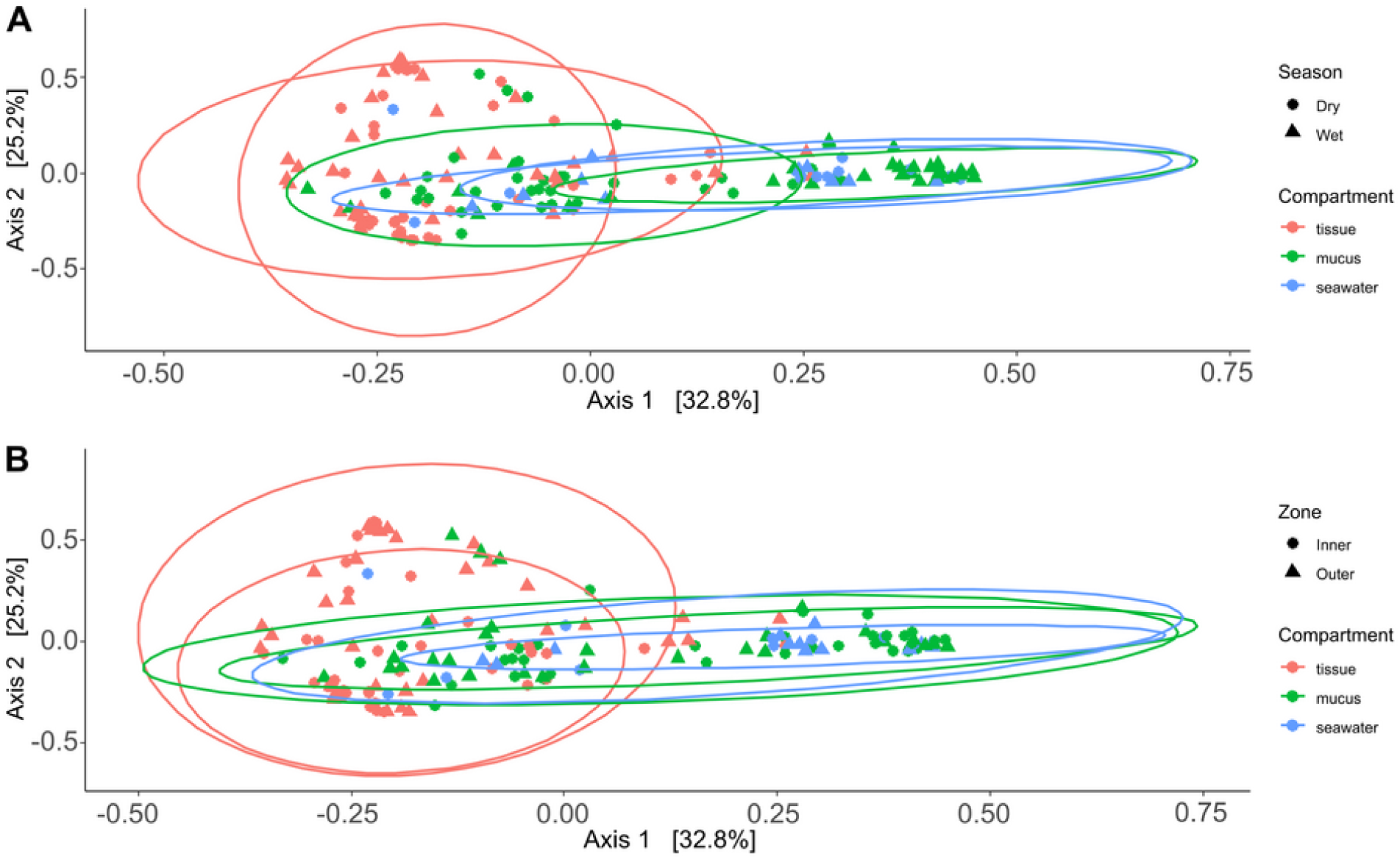
Principal Coordinate Analysis (PCoA) of weighted UniFrac distances. The first two axes of a PCoA of weighted UniFrac distances were plotted to visualize beta diversity patterns, showing that tissue microbiomes were distinct but overlapping with mucus and seawater microbiomes. The contribution of each axis to overall variation in the dataset is provided in percent. Shapes (circles and triangles) were used to highlight possible beta diversity patterns across seasons (A) and reef zones (B).

Pairwise PERMANOVAs (Supplemental Table 6) revealed significant differences in beta diversity between each of the different compartments. Overall, significant differences were also seen between months: April and December, and July and September. No significant monthly differences were seen within the tissue microbiome with the exception of September and December. In mucus samples, significant differences were seen between each month except for September and December. Seawater samples saw significant differences between April and July as well as September and December alone.

### Compartment-specific microbiome differences

Tissue microbiomes were characterized by Endozoicomonadaceae, Simkaniaceae, Alphaproteobacteria, and Myxococcaceae that were significantly more abundant in tissue microbiomes compared to the mucus (Fig. 7A). In the mucus, Sphingomonadaceae, Comamonadaceae, Moraxellaceae, and Pseudomonadaceae were significantly more abundant than in tissues (Fig. 7A). When comparing the mucus microbiome to seawater, the most striking difference was the increased abundance of Alteromonadaceae in seawater compared to the mucus (Fig. 7B). The most pronounced difference across compartments were the high abundance of Endozoicomonadaceae and Simkaniaceae in tissues, Sphingomonadaceae in the mucus, and Alteromonadaceae in the seawater (Fig. 7).

**Figure 7.**
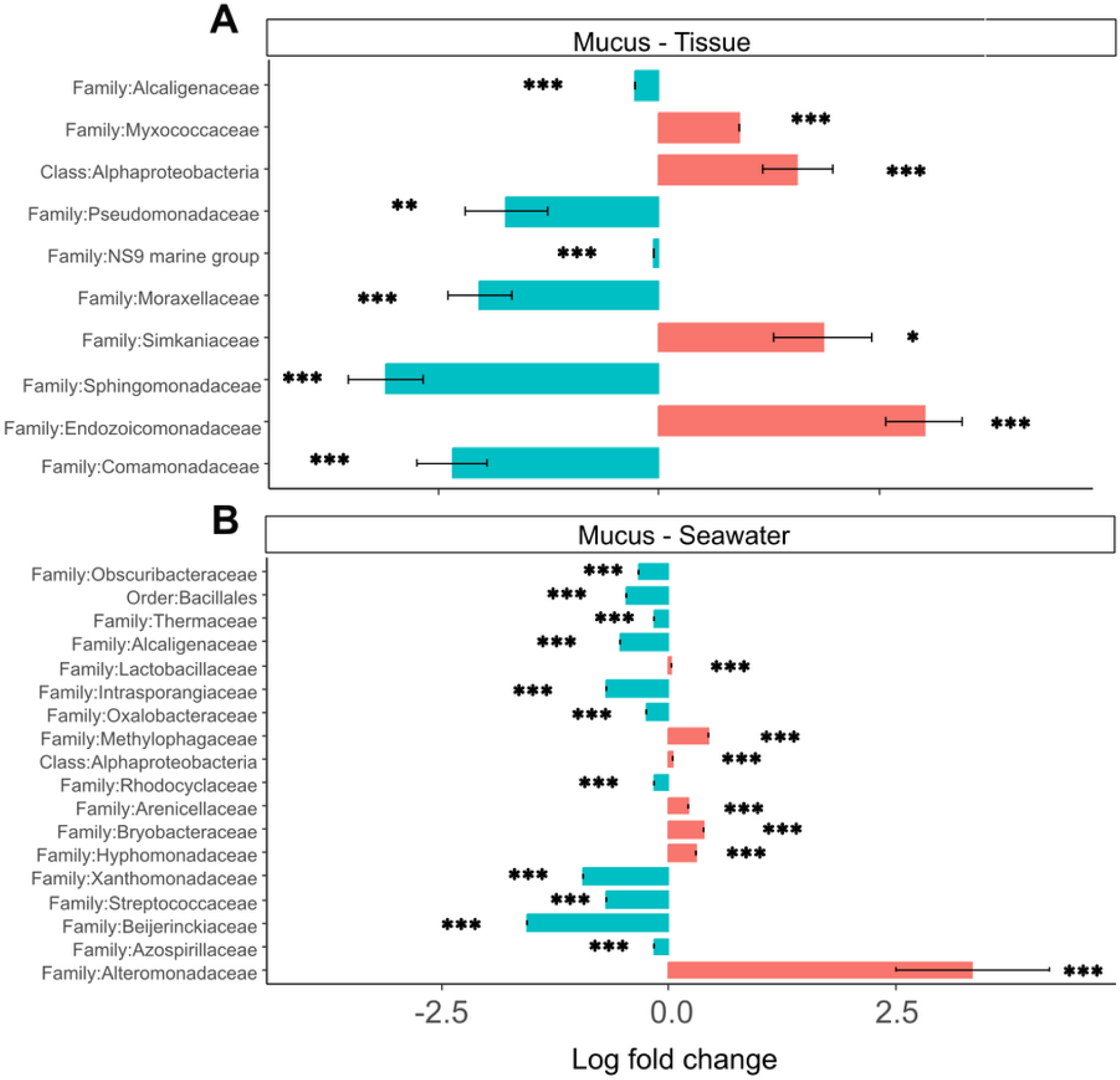
Differential abundance of bacterial taxa in microbiomes. Bacterial taxa that were differentially abundant in comparisons of mucus *versus* tissue and mucus *versus* seawater microbiomes are shown. Asterisks indicate significance levels (*: p ≤ 0.05; **: p ≤ 0.01; ***: p ≤ 0.001).

## Discussion

Bacteria are an essential component of the coral symbiotic community that ensures homeostasis of the holobiont (Boilard et al., 2020). Environmental change and bacterial pollution may lead to disruptions of the coral bacterial microbiome community, which can leave coral vulnerable to stress and disease (Haapkylä et al., 2011). Rainfall patterns during our study period reflected distinct wet and dry seasons with relatively low precipitation during the first two sampling time points in April and at the beginning of July that drastically increased by September, the third sampling time point (Supplemental Fig. 2). While tissue microbiome taxonomic composition remained relatively stable throughout the year (Fig. 5A), mucus microbiome taxonomic composition showed a distinct shift between the dry (April and July) and wet (September and and December) seasons (Fig. 5B). As may be expected, seawater microbiomes appeared most variable, differing across all time points, albeit sample sizes were rather limited (Fig. 5C).

### Potential drivers of microbiome diversity

Consistent with rainfall patterns and resulting runoff into West Hagåtña Bay, *E. coli* and total coliform concentrations increased from July to September and remained high throughout December (Figs. 2B & C); high concentrations of coliform bacteria, including *E. coli* (Figs. 2B & C), in April were the likely result of a rain event the day prior to sampling. *Enterococcus* concentrations were elevated from July until the end of the wet season in December (Fig. 2A). While enterococci are used as indicators of sewage pollution similar to coliform bacteria, *Enterococcus* species may grow in different environments making them more general indicators of non-point source pollution from reservoirs including, for example, soils in addition to sewage (Rothenheber and Jones, 2018). Considering that culverts drain residential runoff directly into the inner zone of West Hagåtña Bay, it is not surprising that FIB concentrations were elevated in the inner zone compared to the outer zone in September during the height of the wet season (Fig. 2B).

We found rainfall to be positively associated with FIB concentrations, in particular coliform and *E. coli* concentrations (Fig. 4) that in turn were identified as weak but significant predictors of tissue and mucus microbiome diversity (Fig. 4). Water temperatures were negatively correlated with FIB concentrations and differed between inner and outer zones (Supplemental Fig. 1). The SEM (Fig. 4) identified seasonal rainfall patterns and associated FIB concentrations as drivers of microbiome diversity. Depending on the metric used, variations in both tissue and mucus microbiome diversity were explained by month or the interaction of zone and month (Supplemental Tables 2-4). Note that zone alone was not able to explain differences in microbiome diversity, which suggests that the combination of increased runoff caused by seasonal rainfalls and distance from shore (inner *versus* outer zone) influenced microbiome diversity in *A. pulchra*.

### Spatiotemporal patterns of microbiome composition

Previous studies compared coral microbiomes across different compartments (Pollock et al., 2018; Marchioro et al., 2020; Sweet, Croquer and Bythell, 2011), microhabitats (Camp et al., 2020; Fifer at al., 2022), or examined microbiome shifts over time (Dunphy et al., 2019; Chu and Vollmer, 2016; Sweet, Croquer and Bythell, 2010). In this study, we tracked the microbiome of *A. pulchra*’s tissue and mucus over time, comparing two habitats, near-shore inner and far-shore outer zones of West Hagåtña Bay. The inner zone of West Hagåtña Bay experienced higher water temperatures (Supplemental Fig. 1) compared to the outer zone. *A. pulchra* has previously been shown to bleach more severely in the inner zone compared to its conspecifics in the outer zone (Raymundo et al., 2017), likely caused by a combination of water temperature and flow differences (Fifer et al., 2021). We uncovered high relative abundances of Simkaniaceae and Endozoicomonadaceae in the tissues of inner zone *A. pulchra* (Fig, 5A), taxa that were characteristic of tissue microbiomes (Fig. 7A). Together, these two bacterial taxa can act as an important energy source for their coral host (Maire et al., 2023). Given that coral bleaching disrupts the nutritional symbiosis between corals and their algal Symbiodiniaceae endosymbionts (Boilard et al., 2020), the high relative abundance of Simkaniaceae suggests microbiome acclimation to the environment of the inner zone of West Hagåtña Bay in *A. pulchra*. While recent years have seen increasing research on the role of Endozoicomonaceae on coral holobiont function, the potential importance of Simkaniaceae for coral holobiont health has only been recognized recently (Maire et al., 2023).

Tissue microbiomes were dominated by Endozoicomonaceae and Simkaniaceae throughout the year, but mucus microbiomes displayed dramatic shifts in taxonomic composition between dry (April and July) and wet (September and December) seasons (Fig. 5B). These seasonal differences in taxonomic composition were also reflected in microbiome diversity that varied between month of sampling and zone (Supplemental Tables 2-4). The high variability and increasing diversity of mucus microbiomes during periods of high rainfall suggests that the mucus bacterial community is impacted by or responding to environmental change. While mucus bacterial communities were dominated by Endozoicomonaceae during April and July (Fig. 5B), similar to tissue microbiomes (Fig. 5A), relative abundances of Endozoicomonaceae in the mucus declined dramatically during the wet season in September and December.

Runoff may increase total dissolved solids, particulate nutrients that are frequently elevated in Guam’s wet season (Guam Waterworks Authority, 2019), which may have led to the growth of the abundant Rhodobacteraceae and Cyanobiaceae present in seawater in September (Fig. 5C). Several mucus samples included high relative abundances for these taxa as well (Fig. 5B), likely originating from seawater. By December, Comamonadaceae, Moraxellaceae, Chitinophagaceae, and Pseudomonadaceae were the most abundant mucus microbiome taxa with community compositions relatively homogenous across samples (Fig. 5B), suggesting microbiome acclimation. While their function remains largely unknown, these taxa have been found in the microbiomes of healthy corals (Chu and Vollmer, 2016; McKew et al., 2012; Vijay et al., 2021) and may provide benefits for *A. pulchra* when exposed to increased runoff during the wet season given their overrepresentation in the mucus (Fig. 7A).

Our initial expectation was that seawater bacterial communities would resemble mucus microbiomes given their direct contact. Indeed, both Shannon diversity and evenness of seawater and mucus bacterial communities were more similar to each other and higher than those of tissue microbiomes (Figs. 3C & D). For observed and phylogenetic diversity, the pattern was opposite (Figs. 3A & B), indicating that the mucus contains fewer bacterial taxa than seawater. However, taxonomic composition of microbiome communities is more similar between the mucus and seawater compared to tissues (Fig. 5). While microbiome composition does not differ across zones for any compartment (Fig. 6B), season does exert some influence on beta diversity with dry season mucus microbiomes showing more overlap with tissue microbiomes than wet season mucus microbiomes (Fig. 6A). Consistent with these observations and the seasonal variations of mucus and seawater microbiome composition (Figs. 5B & C), sampling month was the sole factor explaining variation in the bacterial species pool of these compartments (Supplemental Table 5). By contrast, tissue microbiome composition was affected by zone and the interaction of zone and sampling month (Supplemental Table 5), the likely result of the relatively high abundance of Simkaniaceae during several months, particularly in the inner zone (Fig. 5A).

### Microbiome conformism and regulation

Generally, *Acropora* spp. are considered conformers whose microbiomes are susceptible to and reflect shifts in the external environment (Ziegler et al., 2019). However, some acroporids have been shown to be capable of microbiome regulation. For example, *A. tenuis* is highly susceptible to the influence of its microbiome by the external environment while *A. millepora* has the ability to regulate its microbiome, dampening the impact of seasonal environmental changes (Marchioro et al., 2020). When challenged with a new environment following transplantation, *A. hyacinthus* microbiomes appear to acclimate to new and stressful conditions (Ziegler et al., 2017). In *A. pulchra*, we found mucus and seawater bacterial communities to be similar (Fig. 6), suggesting that the mucus conforms to its surrounding environment. Tissue microbiomes, on the other hand, were relatively distinct (Fig. 6) and stable through time with community shifts largely restricted to changes in relative abundances of the dominant taxa Endozoicomonadaceae and Simkaniaceae (Fig. 5A). Consistent with previous studies (Marchioro et al., 2020; Sweet, Croquer and Bythell, 2011; Apprill, Weber and Santoro, 2016), we also observed tissue microbiomes to be less diverse compared to mucus and seawater bacterial communities (Fig. 3).

Bulk microbiomes are often extracted from combined tissue and mucus samples with the explicit aim of characterizing microbiomes of the entire coral holobiont (e.g., Pootakham et al., 2021). Considering that environmental influences are not uniform across the different microbiome compartments, examining microbiomes of the different layers of the coral holobiont separately has the potential to provide important insights into coral holobiont responses to environmental impacts. In *A. pulchra*, we found that the mucus acts as a microbiome conformer while the stability and relatively low diversity of tissue microbiomes suggests at least some degree of regulation. *A. pulchra* tissues were dominated by Endozoicomonadaceae, which are considered essential coral endosymbionts that produce antimicrobial compounds (Rua et al., 2014) and are positively correlated with Symbiodiniaceae densities (Marchioro et al., 2020). Relative abundance of Simkaniaceae appears highest in the inner zone, near the shore (Fig. 5A).

Consistent with the results presented here, colonies of *A. pulchra* growing near the reef crest in New Caledonia were dominated by *Endozoicomonas* while those living in lagoons were dominated by Simkaniaceae but also Moraxellaceae (Camp et al., 2020). Simkaniaceae have been documented as dominant constituents of juvenile coral microbiomes and are thought to be obligate intracellular endosymbionts (Vouga et al., 2017). Simkaniaceae have been found in cell-associated microbial aggregates (CAMAs) co-occurring with *Endozoicomonas* (Maire et al., 2023). *Endozoicomonas* spp. may secrete excess acetate which *Simkania* spp. may use as an energy source, ultimately aiding coral metabolism (Maire et al., 2023). The increased abundance of Simkaniaceae in inner zone *A. pulchra* tissues but also the outer zone in September during the height of the wet season may represent an acclimation response that aids *A. pulchra* in mitigating the impacts of elevated water temperatures and rainfall-driven coastal runoff.

Mucus microbiomes appeared to have been disrupted early in the wet season, with relative abundances of Endozoicomonaceae declining (Fig. 5B). Interestingly, Rhodobacteraceae were highly abundant in seawater and some mucus bacterial communities in September (Figs. 5B & C). Rhodobacteraceae co-occured with Cyanobiaceae, both of which are commonly found together (Deignan and McDougald, 2022; Botte et al., 2022).

Rhodobacteraceae is a family comprising opportunistic heterotrophic bacteria that rapidly grow in the presence of organic-rich matter, as may be supplied by terrestrial runoff (McDevitt-Irwin et al., 2017).

High relative abundance of Rhodobacteraceae have been associated with declining Endozoicomonadaceae abundance, suggesting negative coral health impacts (Pootakham et al., 2019). By December, Rhodobacteraceae abundances had declined in mucus microbiomes and seawater (Figs. 5B & C) with mucus mirobiomes dominated by Comamonadaceae, Moraxellaceae, Chitinophagaceae, and Pseudomonadaceae. Pseudomonadaceae have previously been described from the mucus of healthy corals and are known for their antibacterial, antiviral, and antifouling properties that allow for control of viruses and prevention of biofilm formation (Vijay et al., 2021), linking this taxon to potential roles in microbiome regulation. Moraxellaceae, such as *Psychrobacter* spp., have also been described from coral mucus samples (McKew et al., 2012) and possess genes associated with carbon and nitrogen metabolism that may confer the ability to utilize organic compounds found in the mucus (Badhai, Ghosh and Das, 2016). While their functional role is not well understood, Comamonadaceae has been reported from healthy corals and macroalgae found in reefs not impacted by environmental stressors (Chu and Vollmer, 2016; Barott et al., 2011; Roder et al., 2014).

Despite being in direct contact with the environment, mucus microbiome composition may in part be regulated by the coral host’s physiology (Glasl, Herndle and Frade, 2016). *A. pulchra* mucus microbiomes were highly variable in September and drastically differed from mucus microbiomes sampled during earlier time points (Fig. 5B). By December, mucus microbiomes were relatively uniform but different from the microbiomes dominated largely by Endozoicomonadaceae in the dry season (Fig. 5B). This pattern suggests that mucus microbiomes were disrupted by environmental changes in the wet season, followed by possible coral host regulation of microbiome composition to a new state characterized by beneficial bacterial taxa.

In addition, Alteromonadaceae were abundant in seawater during September (Fig. 5C) and were generally overrepresented in seawater compared to the mucus (Fig. 7B). Alteromonadaceae such as *Alteromonas* live freely in seawater and are associated with incorporation and possible translocation of nutrients to corals (Ceh et al., 2013). While we found no evidence of elevated concentrations of dissolved nitrogen or phosphate contrary to the results of FIB monitoring (Fig. 3), signatures of long-term eutrophication (Redding et al., 2013) as well as regular detection of elevated levels of total suspended solids (TSS) have been reported from West Hagåtña Bay (Guam Waterworks Authority, 2019).

## Conclusions

This study characterized the bacterial microbiome of coral tissue and mucus of the staghorn coral *Acropora pulchra* growing in nearshore (close to the shore) and farshore (close to the reef crest) habitats to examine shifts in microbiome diversity and composition between wet and dry seasons. Microbiome diversity in both coral tissues and mucus was influenced by seasonal rainfall patterns and associated bacterial pollution, highlighting the impact of runoff on coral microbiomes. However, *Acropora pulchra* tissue microbiomes remained relatively stable across space and time despite *Acropora* species being considered microbiome conformers whose microbiomes closely resemble the surrounding environment. By contrast, *Acropora pulchra* mucus microbiomes were highly variable, with distinct differences in microbiome composition associated with the transition from dry to wet season. This study highlights the differing effects of coastal runoff and bacterial pollution on different compartments of the coral bacterial microbiome and potential drivers of coral microbiome diversity.

## Supporting information

Data analysis scripts

## Acknowledgments

We would like to thank all who contributed to this project, particularly Justin Berg, who assisted with field work and sampling. TCM would like to acknowledge Dr. Laurie Raymundo and Dr. Rebecca Vega Thurber for serving on her MS thesis committee on which the work presented here is based. We would like to thank Dr. Héloïse Rouzé for assisting with bioinformatic data analysis, particularly for her instructions on the use of DADA2 and phyloseq. We would further like to thank Dr. Brett Taylor for feedback on the construction of the structural equation model presented herein.

## Funding

This work was supported by the National Science Foundation under cooperative agreement OIA-1946352. Any opinions, findings, conclusions, or recommendations expressed in this material are those of the authors and do not necessarily reflect the views of the National Science Foundation.

## Notes

### Competing Interest Statement

The authors have declared no competing interest.

