## Supplemental Material for "Seasonal dynamics and environmental drivers of tissue and mucus microbiomes in the staghorn coral *Acropora pulchra*"

**Supplemental Table 1.** GenBank accession numbers for metabarcoding data included in the analyses. All samples were submitted to GenBank under BioProject PRJNA1011454. Collection dates and coordinates are provided for each sample. The tag number represents the unique tag used to identify coral colonies that were repeatedly sampled during the course of the study. Non-chimeric sequences identified by DADA2 are provided for each sample. SRA: Sequence Read Archive.

| Biosample Accession | SRA Accession | Collection Date | Coordinates | Tag # | Sample Type | Non-chimeric Sequences |
|---|---|---|---|---|---|---|
| SAMN37218957 | SRR25867451 | 2021-04-24 | 13.482 N 144.746 E | 61 | Tissue | 37,952 |
| SAMN37219041 | SRR25867428 | 2021-04-24 | 13.482 N 144.745 E | 61 | Mucus | 250,740 |
| SAMN37218970 | SRR25867350 | 2021-07-01 | 13.482 N 144.746 E | 61 | Tissue | 292,933 |
| SAMN37219054 | SRR25867348 | 2021-07-01 | 13.482 N 144.746 E | 61 | Mucus | 6,091 |
| SAMN37218993 | SRR25867293 | 2021-09-21 | 13.482 N 144.746 E | 61 | Tissue | 49,361 |
| SAMN37219077 | SRR25867292 | 2021-09-21 | 13.482 N 144.746 E | 61 | Mucus | 20,527 |
| SAMN37219015 | SRR25867435 | 2021-12-28 | 13.482 N 144.746 E | 61 | Tissue | 249,543 |
| SAMN37219099 | SRR25867431 | 2021-12-28 | 13.482 N 144.746 E | 61 | Mucus | 8,838 |
| SAMN37218955 | SRR25867301 | 2021-04-24 | 13.482 N 144.746 E | 62 | Tissue | 237,022 |
| SAMN37219039 | SRR25867381 | 2021-04-24 | 13.482 N 144.745 E | 62 | Mucus | 158,651 |
| SAMN37218971 | SRR25867345 | 2021-07-01 | 13.482 N 144.746 E | 62 | Tissue | 216,788 |
| SAMN37219055 | SRR25867344 | 2021-07-01 | 13.482 N 144.746 E | 62 | Mucus | 28,341 |
| SAMN37218994 | SRR25867291 | 2021-09-21 | 13.482 N 144.746 E | 62 | Tissue | 2,764 |
| SAMN37219078 | SRR25867290 | 2021-09-21 | 13.482 N 144.746 E | 62 | Mucus | 111,027 |
| SAMN37219016 | SRR25867433 | 2021-12-28 | 13.482 N 144.746 E | 62 | Tissue | 122,195 |
| SAMN37219100 | SRR25867429 | 2021-12-28 | 13.482 N 144.746 E | 62 | Mucus | 51,753 |
| SAMN37218958 | SRR25867427 | 2021-04-24 | 13.482 N 144.746 E | 63 | Tissue | 166,360 |
| SAMN37219042 | SRR25867409 | 2021-04-24 | 13.482 N 144.744 E | 63 | Mucus | 141,616 |
| SAMN37218972 | SRR25867343 | 2021-07-01 | 13.482 N 144.746 E | 63 | Tissue | 13,377 |
| SAMN37219056 | SRR25867342 | 2021-07-01 | 13.482 N 144.746 E | 63 | Mucus | 9,123 |
| SAMN37218995 | SRR25867389 | 2021-09-21 | 13.482 N 144.746 E | 63 | Tissue | 40,208 |
| SAMN37219079 | SRR25867388 | 2021-09-21 | 13.482 N 144.746 E | 63 | Mucus | 12,034 |
| SAMN37219017 | SRR25867430 | 2021-12-28 | 13.482 N 144.746 E | 63 | Tissue | 107,183 |
| SAMN37219101 | SRR25867425 | 2021-12-28 | 13.482 N 144.745 E | 63 | Mucus | 93,048 |
| SAMN37218952 | SRR25867456 | 2021-04-24 | 13.482 N 144.745 E | 64 | Tissue | 37,509 |
| SAMN37219036 | SRR25867347 | 2021-04-24 | 13.482 N 144.746 E | 64 | Mucus | 145,403 |
| SAMN37218974 | SRR25867339 | 2021-07-01 | 13.482 N 144.745 E | 64 | Tissue | 271,399 |
| SAMN37218996 | SRR25867387 | 2021-09-21 | 13.482 N 144.745 E | 64 | Tissue | 45,210 |
| SAMN37219080 | SRR25867386 | 2021-09-21 | 13.482 N 144.745 E | 64 | Mucus | 105,101 |
| SAMN37219018 | SRR25867426 | 2021-12-28 | 13.482 N 144.745 E | 64 | Tissue | 64,736 |
| SAMN37219102 | SRR25867423 | 2021-12-28 | 13.482 N 144.745 E | 64 | Mucus | 14,999 |
| SAMN37218960 | SRR25867356 | 2021-04-24 | 13.482 N 144.745 E | 65 | Tissue | 43,569 |
| SAMN37219044 | SRR25867353 | 2021-04-24 | 13.482 N 144.744 E | 65 | Mucus | 32,839 |
| SAMN37218975 | SRR25867336 | 2021-07-01 | 13.482 N 144.745 E | 65 | Tissue | 170,389 |
| SAMN37219059 | SRR25867335 | 2021-07-01 | 13.482 N 144.745 E | 65 | Mucus | 89,345 |
| SAMN37218997 | SRR25867385 | 2021-09-21 | 13.482 N 144.745 E | 65 | Tissue | 68,876 |
| SAMN37219081 | SRR25867384 | 2021-09-21 | 13.482 N 144.745 E | 65 | Mucus | 38,654 |

| | | | | | | |
|---|---|---|---|---|---|---|
| SAMN37219019 | SRR25867424 | 2021-12-28 | 13.482 N 144.745 E | 65 | Tissue | 23,570 |
| SAMN37219103 | SRR25867422 | 2021-12-28 | 13.482 N 144.745 E | 65 | Mucus | 19,789 |
| SAMN37218954 | SRR25867324 | 2021-04-24 | 13.482 N 144.745 E | 66 | Tissue | 64,089 |
| SAMN37219038 | SRR25867302 | 2021-04-24 | 13.482 N 144.746 E | 66 | Mucus | 205,286 |
| SAMN37218976 | SRR25867334 | 2021-07-01 | 13.482 N 144.745 E | 66 | Tissue | 135,976 |
| SAMN37219060 | SRR25867333 | 2021-07-01 | 13.482 N 144.745 E | 66 | Mucus | 4,400 |
| SAMN37218998 | SRR25867382 | 2021-09-21 | 13.482 N 144.745 E | 66 | Tissue | 76,535 |
| SAMN37218956 | SRR25867380 | 2021-04-24 | 13.482 N 144.744 E | 67 | Tissue | 115,382 |
| SAMN37219040 | SRR25867452 | 2021-04-24 | 13.482 N 144.745 E | 67 | Mucus | 128,529 |
| SAMN37218978 | SRR25867330 | 2021-07-01 | 13.482 N 144.744 E | 67 | Tissue | 92,596 |
| SAMN37219062 | SRR25867329 | 2021-07-01 | 13.482 N 144.744 E | 67 | Mucus | 10,118 |
| SAMN37219000 | SRR25867377 | 2021-09-21 | 13.482 N 144.744 E | 67 | Tissue | 114,596 |
| SAMN37219084 | SRR25867374 | 2021-09-21 | 13.482 N 144.744 E | 67 | Mucus | 58,916 |
| SAMN37219021 | SRR25867418 | 2021-12-28 | 13.482 N 144.744 E | 67 | Tissue | 25,884 |
| SAMN37219105 | SRR25867417 | 2021-12-28 | 13.482 N 144.744 E | 67 | Mucus | 94,909 |
| SAMN37218953 | SRR25867346 | 2021-04-24 | 13.482 N 144.744 E | 68 | Tissue | 62,026 |
| SAMN37218981 | SRR25867323 | 2021-07-01 | 13.482 N 144.744 E | 68 | Tissue | 198,967 |
| SAMN37219065 | SRR25867320 | 2021-07-01 | 13.480 N 144.743 E | 68 | Mucus | 1,440 |
| SAMN37219001 | SRR25867375 | 2021-09-21 | 13.482 N 144.744 E | 68 | Tissue | 308,652 |
| SAMN37219085 | SRR25867371 | 2021-09-21 | 13.482 N 144.744 E | 68 | Mucus | 49,694 |
| SAMN37219022 | SRR25867416 | 2021-12-28 | 13.482 N 144.744 E | 68 | Tissue | 21,117 |
| SAMN37219106 | SRR25867415 | 2021-12-28 | 13.482 N 144.744 E | 68 | Mucus | 30,154 |
| SAMN37218961 | SRR25867455 | 2021-04-24 | 13.482 N 144.744 E | 69 | Tissue | 57,519 |
| SAMN37219045 | SRR25867349 | 2021-04-24 | 13.480 N 144.743 E | 69 | Mucus | 99,525 |
| SAMN37218980 | SRR25867326 | 2021-07-01 | 13.482 N 144.744 E | 69 | Tissue | 51,353 |
| SAMN37219064 | SRR25867322 | 2021-07-01 | 13.482 N 144.744 E | 69 | Mucus | 66,907 |
| SAMN37219002 | SRR25867373 | 2021-09-21 | 13.482 N 144.744 E | 69 | Tissue | 70,306 |
| SAMN37219086 | SRR25867369 | 2021-09-21 | 13.480 N 144.743 E | 69 | Mucus | 45,020 |
| SAMN37219023 | SRR25867414 | 2021-12-28 | 13.482 N 144.744 E | 69 | Tissue | 63,130 |
| SAMN37219107 | SRR25867413 | 2021-12-28 | 13.482 N 144.744 E | 69 | Mucus | 82,328 |
| SAMN37218967 | SRR25867421 | 2021-04-24 | 13.480 N 144.743 E | 72 | Tissue | 133,786 |
| SAMN37219051 | SRR25867410 | 2021-04-24 | 13.480 N 144.743 E | 72 | Mucus | 108,955 |
| SAMN37218989 | SRR25867304 | 2021-07-01 | 13.480 N 144.743 E | 72 | Tissue | 22,657 |
| SAMN37219011 | SRR25867444 | 2021-09-21 | 13.480 N 144.743 E | 72 | Tissue | 55,748 |
| SAMN37219095 | SRR25867440 | 2021-09-21 | 13.480 N 144.743 E | 72 | Mucus | 21,057 |
| SAMN37219032 | SRR25867392 | 2021-12-28 | 13.480 N 144.743 E | 72 | Tissue | 14,659 |
| SAMN37219116 | SRR25867391 | 2021-12-28 | 13.480 N 144.743 E | 72 | Mucus | 80,635 |
| SAMN37218968 | SRR25867399 | 2021-04-24 | 13.480 N 144.743 E | 73 | Tissue | 167,723 |
| SAMN37219052 | SRR25867360 | 2021-04-24 | 13.480 N 144.743 E | 73 | Mucus | 25,735 |
| SAMN37218990 | SRR25867300 | 2021-07-01 | 13.480 N 144.743 E | 73 | Tissue | 41,517 |
| SAMN37219074 | SRR25867299 | 2021-07-01 | 13.480 N 144.743 E | 73 | Mucus | 1,023 |
| SAMN37219012 | SRR25867441 | 2021-09-21 | 13.480 N 144.743 E | 73 | Tissue | 105,278 |
| SAMN37219096 | SRR25867438 | 2021-09-21 | 13.480 N 144.743 E | 73 | Mucus | 38,448 |
| SAMN37219033 | SRR25867390 | 2021-12-28 | 13.480 N 144.743 E | 73 | Tissue | 30,351 |
| SAMN37219117 | SRR25867362 | 2021-12-28 | 13.480 N 144.743 E | 73 | Mucus | 29,716 |
| SAMN37218969 | SRR25867352 | 2021-04-24 | 13.480 N 144.743 E | 74 | Tissue | 126,345 |
| SAMN37219053 | SRR25867351 | 2021-04-24 | 13.480 N 144.743 E | 74 | Mucus | 17,136 |
| SAMN37218991 | SRR25867298 | 2021-07-01 | 13.480 N 144.743 E | 74 | Tissue | 43,890 |

| | | | | | | |
|---|---|---|---|---|---|---|
| SAMN37219075 | SRR25867297 | 2021-07-01 | 13.480 N 144.743 E | 74 | Mucus | 1,365 |
| SAMN37219013 | SRR25867439 | 2021-09-21 | 13.480 N 144.743 E | 74 | Tissue | 76,272 |
| SAMN37219034 | SRR25867359 | 2021-12-28 | 13.480 N 144.743 E | 74 | Tissue | 31,642 |
| SAMN37219118 | SRR25867358 | 2021-12-28 | 13.480 N 144.743 E | 74 | Mucus | 111,241 |
| SAMN37218964 | SRR25867294 | 2021-04-24 | 13.480 N 144.743 E | 75 | Tissue | 125,093 |
| SAMN37219048 | SRR25867383 | 2021-04-24 | 13.480 N 144.743 E | 75 | Mucus | 8,982 |
| SAMN37218985 | SRR25867313 | 2021-07-01 | 13.480 N 144.743 E | 75 | Tissue | 362,795 |
| SAMN37219069 | SRR25867312 | 2021-07-01 | 13.480 N 144.743 E | 75 | Mucus | 20,478 |
| SAMN37219007 | SRR25867361 | 2021-09-21 | 13.480 N 144.743 E | 75 | Tissue | 43,557 |
| SAMN37219091 | SRR25867449 | 2021-09-21 | 13.480 N 144.743 E | 75 | Mucus | 97,566 |
| SAMN37219028 | SRR25867401 | 2021-12-28 | 13.480 N 144.743 E | 75 | Tissue | 15,022 |
| SAMN37219112 | SRR25867400 | 2021-12-28 | 13.480 N 144.743 E | 75 | Mucus | 18,200 |
| SAMN37218965 | SRR25867372 | 2021-04-24 | 13.480 N 144.743 E | 76 | Tissue | 82,674 |
| SAMN37219049 | SRR25867454 | 2021-04-24 | 13.480 N 144.743 E | 76 | Mucus | 91,676 |
| SAMN37218986 | SRR25867311 | 2021-07-01 | 13.480 N 144.743 E | 76 | Tissue | 95,065 |
| SAMN37219070 | SRR25867310 | 2021-07-01 | 13.480 N 144.743 E | 76 | Mucus | 1,641 |
| SAMN37219008 | SRR25867450 | 2021-09-21 | 13.480 N 144.743 E | 76 | Tissue | 193,058 |
| SAMN37219092 | SRR25867447 | 2021-09-21 | 13.480 N 144.743 E | 76 | Mucus | 99,413 |
| SAMN37219029 | SRR25867398 | 2021-12-28 | 13.480 N 144.743 E | 76 | Tissue | 20,489 |
| SAMN37219113 | SRR25867397 | 2021-12-28 | 13.480 N 144.743 E | 76 | Mucus | 28,788 |
| SAMN37218966 | SRR25867432 | 2021-04-24 | 13.480 N 144.743 E | 77 | Tissue | 67,824 |
| SAMN37219050 | SRR25867443 | 2021-04-24 | 13.480 N 144.743 E | 77 | Mucus | 274,535 |
| SAMN37218987 | SRR25867309 | 2021-07-01 | 13.480 N 144.743 E | 77 | Tissue | 85,978 |
| SAMN37219071 | SRR25867308 | 2021-07-01 | 13.480 N 144.743 E | 77 | Mucus | 2,408 |
| SAMN37219009 | SRR25867448 | 2021-09-21 | 13.480 N 144.743 E | 77 | Tissue | 45,625 |
| SAMN37219030 | SRR25867396 | 2021-12-28 | 13.480 N 144.743 E | 77 | Tissue | 25,829 |
| SAMN37219114 | SRR25867395 | 2021-12-28 | 13.480 N 144.743 E | 77 | Mucus | 63,828 |
| SAMN37218962 | SRR25867338 | 2021-04-24 | 13.480 N 144.742 E | 78 | Tissue | 62,281 |
| SAMN37219046 | SRR25867327 | 2021-04-24 | 13.480 N 144.743 E | 78 | Mucus | 83,536 |
| SAMN37218982 | SRR25867321 | 2021-07-01 | 13.480 N 144.743 E | 78 | Tissue | 29,620 |
| SAMN37219066 | SRR25867319 | 2021-07-01 | 13.480 N 144.743 E | 78 | Mucus | 1,311 |
| SAMN37219003 | SRR25867370 | 2021-09-21 | 13.480 N 144.743 E | 78 | Tissue | 58,969 |
| SAMN37219087 | SRR25867367 | 2021-09-21 | 13.480 N 144.743 E | 78 | Mucus | 43,578 |
| SAMN37219024 | SRR25867412 | 2021-12-28 | 13.480 N 144.743 E | 78 | Tissue | 20,544 |
| SAMN37219108 | SRR25867411 | 2021-12-28 | 13.480 N 144.743 E | 78 | Mucus | 90,894 |
| SAMN37218963 | SRR25867316 | 2021-04-24 | 13.480 N 144.742 E | 79 | Tissue | 344,464 |
| SAMN37219047 | SRR25867305 | 2021-04-24 | 13.480 N 144.743 E | 79 | Mucus | 562,259 |
| SAMN37219004 | SRR25867368 | 2021-09-21 | 13.480 N 144.743 E | 79 | Tissue | 68,656 |
| SAMN37219088 | SRR25867365 | 2021-09-21 | 13.480 N 144.743 E | 79 | Mucus | 28,484 |
| SAMN37219025 | SRR25867407 | 2021-12-28 | 13.480 N 144.743 E | 79 | Tissue | 44,474 |
| SAMN37219109 | SRR25867406 | 2021-12-28 | 13.480 N 144.743 E | 79 | Mucus | 141,825 |
| SAMN37218959 | SRR25867408 | 2021-04-24 | 13.480 N 144.742 E | 80 | Tissue | 85,968 |
| SAMN37219043 | SRR25867357 | 2021-04-24 | 13.482 N 144.744 E | 80 | Mucus | 117,351 |
| SAMN37218983 | SRR25867318 | 2021-07-01 | 13.480 N 144.743 E | 80 | Tissue | 157,065 |
| SAMN37219067 | SRR25867317 | 2021-07-01 | 13.480 N 144.743 E | 80 | Mucus | 1,497 |
| SAMN37219005 | SRR25867366 | 2021-09-21 | 13.480 N 144.743 E | 80 | Tissue | 5,316 |
| SAMN37219026 | SRR25867405 | 2021-12-28 | 13.480 N 144.743 E | 80 | Tissue | 64,591 |
| SAMN37219110 | SRR25867404 | 2021-12-28 | 13.480 N 144.743 E | 80 | Mucus | 84,938 |

| | | | | | | |
|---|---|---|---|---|---|---|
| SAMN37239985 | SRR25884572 | 2021-04-24 | 13.480 N 144.743 E | N/A | Seawater | 275,093 |
| SAMN37239986 | SRR25884571 | 2021-04-24 | 13.480 N 144.743 E | N/A | Seawater | 53,670 |
| SAMN37239987 | SRR25884584 | 2021-04-24 | 13.480 N 144.743 E | N/A | Seawater | 472,024 |
| SAMN37239988 | SRR25884579 | 2021-04-24 | 13.482 N 144.746 E | N/A | Seawater | 364,150 |
| SAMN37239989 | SRR25884578 | 2021-04-24 | 13.482 N 144.745 E | N/A | Seawater | 169,642 |
| SAMN37239990 | SRR25884577 | 2021-04-24 | 13.482 N 144.744 E | N/A | Seawater | 145,361 |
| SAMN37239992 | SRR25884591 | 2021-07-01 | 13.480 N 144.743 E | N/A | Seawater | 74,097 |
| SAMN37239993 | SRR25884590 | 2021-07-01 | 13.480 N 144.743 E | N/A | Seawater | 6,731 |
| SAMN37239994 | SRR25884589 | 2021-07-01 | 13.482 N 144.746 E | N/A | Seawater | 7,804 |
| SAMN37239995 | SRR25884588 | 2021-07-01 | 13.482 N 144.745 E | N/A | Seawater | 14,397 |
| SAMN37239996 | SRR25884587 | 2021-07-01 | 13.482 N 144.744 E | N/A | Seawater | 8,035 |
| SAMN37239997 | SRR25884586 | 2021-09-21 | 13.480 N 144.743 E | N/A | Seawater | 15,380 |
| SAMN37239998 | SRR25884585 | 2021-09-21 | 13.480 N 144.743 E | N/A | Seawater | 42,093 |
| SAMN37239999 | SRR25884583 | 2021-09-21 | 13.480 N 144.743 E | N/A | Seawater | 11,884 |
| SAMN37240000 | SRR25884582 | 2021-09-21 | 13.482 N 144.746 E | N/A | Seawater | 11,389 |
| SAMN37240001 | SRR25884581 | 2021-09-21 | 13.482 N 144.745 E | N/A | Seawater | 4,159 |
| SAMN37240002 | SRR25884580 | 2021-09-21 | 13.482 N 144.744 E | N/A | Seawater | 5,448 |
| SAMN37240003 | SRR25884576 | 2021-12-28 | 13.480 N 144.743 E | N/A | Seawater | 67,581 |
| SAMN37240004 | SRR25884575 | 2021-12-28 | 13.480 N 144.743 E | N/A | Seawater | 16,166 |
| SAMN37240005 | SRR25884574 | 2021-12-28 | 13.480 N 144.743 E | N/A | Seawater | 26,744 |
| SAMN37240006 | SRR25884573 | 2021-12-28 | 13.482 N 144.746 E | N/A | Seawater | 52,967 |
| SAMN37240007 | SRR25884570 | 2021-12-28 | 13.482 N 144.745 E | N/A | Seawater | 16,952 |
| SAMN37240008 | SRR25884569 | 2021-12-28 | 13.482 N 144.744 E | N/A | Seawater | 48,214 |

**Supplemental Table 2.** Shannon diversity table for interactions among microbial communities from distinct coral compartments (tissue, mucus and seawater), month (April, July, September and December) and zone (in versus out). Diversity values for overall, tissue, and mucus compartments were calculated using a Kruskal-Wallis Chi2 test. Diversity values for seawater were calculated using an analysis of variance (ANOVA). Significant results (*p*(perm) <0.05) are highlighted in bold.

**Overall**

| Source of Variation<br>Interactions | *df* | $X^2$ | *p*(perm) |
|---|---|---|---|
| Compartment | 2 | 63.138 | **< 0.001** |
| Zone | 1 | 1.0288 | 0.310 |
| Month | 3 | 7.6118 | 0.055 |
| Zone:Month | 7 | 9.2215 | 0.237 |
| Compartment:Zone | 5 | 64.776 | **< 0.001** |
| Compartment:Month | 11 | 73.897 | **< 0.001** |
| Compartment:Zone:Month | 23 | 86.204 | **< 0.001** |

**Tissue**

| Source of Variation<br>Interactions | *df* | $X^2$ | *p*(perm) |
|---|---|---|---|
| Zone | 1 | 0.018247 | 0.893 |
| Month | 3 | 6.1154 | 0.106 |
| Zone:Month | 7 | 13.53 | 0.060 |

**Mucus**

| Source of Variation<br>Interactions | *df* | $X^2$ | *p*(perm) |
|---|---|---|---|
| Zone | 1 | 1.2883 | 0.256 |
| Month | 3 | 8.567 | **0.036** |
| Zone:Month | 7 | 14.507 | **0.043** |

**Seawater**

| Source of Variation<br><br>Interactions | *df* | F value | *p*(perm) |
| --- | --- | --- | --- |
| Zone | 1 | 0.087 | 0.771 |
| Month | 1 | 2.967 | 0.100 |
| Zone:Month | 1 | 0.970 | 0.337 |

**Supplemental Table 3.** Microbial evenness table for interactions among microbial communities from distinct coral compartments (tissue, mucus and seawater), month (April, July, September and December) and zone (in versus out). Evenness values for overall, tissue, and mucus compartments were calculated using a Kruskal-Wallis Chi$^2$ test. Evenness values for seawater were calculated using an analysis of variance (ANOVA). Significant results ($p$(perm) <0.05) are highlighted in bold.

**Overall**

| Source of Variation<br>Interactions | *df* | $X^2$ | *p*(perm) |
|---|---|---|---|
| Compartment | 2 | 64.665 | **< 0.001** |
| Zone | 1 | 0.282 | 0.596 |
| Month | 3 | 2.012 | 0.570 |
| Zone:Month | 7 | 4.000 | 0.780 |
| Compartment:Zone | 5 | 65.649 | **< 0.001** |
| Compartment:Month | 11 | 76.174 | **< 0.001** |
| Compartment:Zone:Month | 23 | 84.665 | **< 0.001** |

**Tissue**

| Source of Variation<br>Interactions | *df* | $X^2$ | *p*(perm) |
|---|---|---|---|
| Zone | 1 | 0.004 | 0.949 |
| Month | 3 | 6.768 | 0.080 |
| Zone:Month | 7 | 15.138 | **0.034** |

**Mucus**

| Source of Variation<br>Interactions | *df* | $X^2$ | *p*(perm) |
|---|---|---|---|
| Zone | 1 | 0.005 | 0.946 |
| Month | 3 | 11.970 | **0.008** |
| Zone:Month | 7 | 15.385 | 0.314 |

**Seawater**

| Source of Variation<br>Interactions | *df* | F value | *p*(perm) |
|---|---|---|---|
| Zone | 1 | 2.362 | 0.139 |
| Month | 1 | 0.040 | 0.844 |
| Zone:Month | 7 | 0.671 | 0.694 |

**Supplemental Table 4.** Faith's Phylogenetic Diversity (PD) table for interactions among microbial communities from distinct compartments (seawater, mucus and tissue), month (April, July, September and December) and zone (in versus out). PD values for overall, tissue, and seawater compartments were calculated using a Kruskal-Wallis Chi$^2$ test. Because mucus has a normal distribution, an analysis of variance (ANOVA) was used. Significant results (*p*(perm) <0.05) are highlighted in bold.

**Overall**

| Source of Variation<br>Interactions | *df* | $X^2$ | *p*(perm) |
|---|---|---|---|
| Compartment | 2 | 15.000 | **< 0.001** |
| Zone | 1 | 0.922 | 0.374 |
| Month | 3 | 27.109 | **< 0.001** |
| Zone:Month | 7 | 31.658 | **< 0.001** |
| Compartment:Zone | 5 | 16.057 | **0.002** |
| Compartment:Month | 11 | 61.849 | **< 0.001** |
| Compartment:Zone:Month | 23 | 72.317 | **< 0.001** |

**Tissue**

| Source of Variation<br>Interactions | *df* | $X^2$ | *p*(perm) |
|---|---|---|---|
| Zone | 1 | 0.164 | 0.685 |
| Month | 3 | 11.218 | **0.011** |
| Zone:Month | 7 | 16.622 | **0.020** |

**Mucus**

| Source of Variation<br>Interactions | *df* | F value | *p*(perm) |
|---|---|---|---|
| Zone | 1 | 0.781 | 0.380 |
| Month | 1 | 2.350 | 0.130 |
| Zone:Month | 7 | 8.105 | **< 0.001** |

**Seawater**

| Source of Variation<br><br>Interactions | *df* | $X^2$ | *p*(perm) |
| --- | --- | --- | --- |
| Zone | 1 | 0.136 | 0.712 |
| Month | 3 | 10.822 | **0.012** |
| Zone:Month | 7 | 12.337 | 0.090 |

**Supplemental Table 5.** Permutational multivariate analysis of variance (PERMANOVA) table for interactions among microbial communities from distinct coral compartments (seawater, mucus and tissue), month (April, July, September and December) and zone (in *versus* out). Significant results (*p*(perm) <0.05) are highlighted in bold.

**Overall**

| Source of Variation<br><br>Interactions | *df* | F value | *p(*perm) |
|---|---|---|---|
| Compartment | 2 | 15.035 | **< 0.001** |
| Zone | 1 | 2.198 | 0.064 |
| Month | 1 | 3.870 | **0.009** |
| Zone:Month | 1 | 3.097 | **0.019** |
| Compartment:Zone | 2 | 1.429 | 0.150 |
| Compartment:Month | 2 | 2.900 | **0.004** |
| Compartment:Zone:Month | 2 | 1.031 | 0.418 |

**Tissue**

| Source of Variation<br><br>Interactions | *df* | F value | *p(*perm) |
|---|---|---|---|
| Zone | 1 | 4.943 | **0.002** |
| Month | 1 | 1.418 | 0.244 |
| Zone:Month | 1 | 4.508 | **0.003** |

**Mucus**

| Source of Variation<br><br>Interactions | *df* | F value | *p(*perm) |
|---|---|---|---|
| Zone | 1 | 1.033 | 0.353 |
| Month | 1 | 7.213 | **< 0.001** |
| Zone:Month | 1 | 1.950 | 0.074 |

**Seawater**

| Source of Variation<br><br>Interactions | *df* | F value | *p*(perm) |
|---|---|---|---|
| Zone | 1 | 0.537 | 0.827 |
| Month | 1 | 1.556 | 0.160 |
| Zone:Month | 1 | 0.291 | 0.982 |

**Supplemental Table 6.** Post-hoc permutational multivariate analysis of variance (PERMANOVA) table for pairwise interactions among microbial communities between individual compartments (seawater, coral mucus and coral tissue), and individual months (May, July, September and December). Significant results (*p*(perm) <0.05) are highlighted in bold.

**Overall**

| Source of Variation<br>Interactions | *df* | Pseudo-*F* | *p*(perm) |
|---|---|---|---|
| Tissue vs. Mucus | 1 | 22.586 | **0.003** |
| Tissue vs. Seawater | 1 | 11.190 | **0.003** |
| Mucus vs. Seawater | 1 | 3.499 | **0.012** |
| Apr vs. Jul | 1 | 3.994 | 0.054 |
| Apr vs. Sep | 1 | 2.163 | 0.234 |
| Apr vs. Dec | 1 | 4.409 | **0.018** |
| Jul vs. Sep | 1 | 4.377 | **0.006** |
| Jul vs. Dec | 1 | 2.731 | 0.168 |
| Sep vs. Dec | 1 | 2.428 | 0.210 |

**Tissue**

| Source of Variation<br>Interactions | *df* | Pseudo-*F* | *p*(perm) |
|---|---|---|---|
| Apr vs. Jul | 1 | 1.464 | 1.000 |
| Apr vs. Sep | 1 | 0.667 | 1.000 |
| Apr vs. Dec | 1 | 2.626 | 0.210 |
| Jul vs. Sep | 1 | 2.806 | 0.132 |
| Jul vs. Dec | 1 | 0.551 | 1.000 |
| Sep vs. Dec | 1 | 4.384 | **0.006** |

**Mucus**

| Source of Variation | *df* | Pseudo-*F* | *p*(perm) |
|---|---|---|---|

| Interactions | | | |
|---|---|---|---|
| Apr vs. Jul | 1 | 11.594 | **0.006** |
| Apr vs. Sep | 1 | 3.299 | **0.024** |
| Apr vs. Dec | 1 | 6.010 | **0.006** |
| Jul vs. Sep | 1 | 9.608 | **0.006** |
| Jul vs. Dec | 1 | 16.631 | **0.006** |
| Sep vs. Dec | 1 | 0.822 | 1.000 |

**Seawater**

| Source of Variation<br>Interactions | *df* | F value | *p*(perm) |
|---|---|---|---|
| Apr vs. Jul | 1 | 2.930 | 0.186 |
| Apr vs. Sep | 1 | 2.279 | 0.713 |
| Apr vs. Dec | 1 | 2.693 | 0.318 |
| Jul vs. Sep | 1 | 3.605 | **0.036** |
| Jul vs. Dec | 1 | 3.098 | 0.126 |
| Sep vs. Dec | 1 | 5.890 | **0.042** |

**Supplemental Table 7**. Phylum-level abundances of bacterial groups in microbiomes sampled. Relative abundances of the six most abundant phyla found in coral tissue, coral mucus, and seawater. Proteobacteria represented the by far most abundant constituents of all microbiomes, in particular coral tissue microbiomes.

| Phylum | Overall % | Tissue % | Mucus % | Seawater % |
|---|---|---|---|---|
| Actinobacteriota | 2.6 | 1.1 | 4.5 | 2.3 |
| Bacteroidota | 5.9 | 1.6 | 6.7 | 19.3 |
| Cyanobacteria | 2.0 | 0.4 | 2.2 | 7.3 |
| Firmicutes | 2.3 | 1.2 | 3.9 | 1.6 |
| Proteobacteria | 76.5 | 83.7 | 72.4 | 63.3 |
| Verrucomicrobiota | 9.9 | 11.7 | 9.3 | 5.2 |

**SupplementalTable 8**. Family-level abundances of bacterial groups in microbiomes sampled. Relative abundances of the 11 most abundant families found overall across all compartments, in coral tissue, in coral mucus, and in seawater are provided. Endozoicomonadaceae were most abundant in all compartments, in particular coral tissue samples.

| Family | Overall % | Tissue % | Mucus % | Seawater % |
|---|---|---|---|---|
| Alteromonadaceae | 1.4 | 0.5 | 1.6 | 4.2 |
| Chitinophageacae | 3.2 | 0.9 | 5.0 | 6.0 |
| Comamonadaceae | 6.8 | 1.6 | 13.6 | 5.1 |
| Cryomorphaceae | 0.8 | 0.1 | 0.4 | 4.7 |
| Cyanobiaceae | 1.7 | 0.3 | 1.6 | 7.3 |
| Endozoicomonadaceae | 51.0 | 76.2 | 31.1 | 23.8 |
| Moraxellaceae | 6.2 | 2.5 | 11.3 | 4.3 |
| Pseudomonadaceae | 1.1 | 0.2 | 2.5 | 0.5 |
| Rhodobacteraceae | 1.3 | 0.1 | 1.2 | 6.3 |
| Simkaniaceae | 9.8 | 11.8 | 9.1 | 4.8 |
| Sphingomonadaceae | 2.1 | 0.5 | 3.8 | 2.8 |

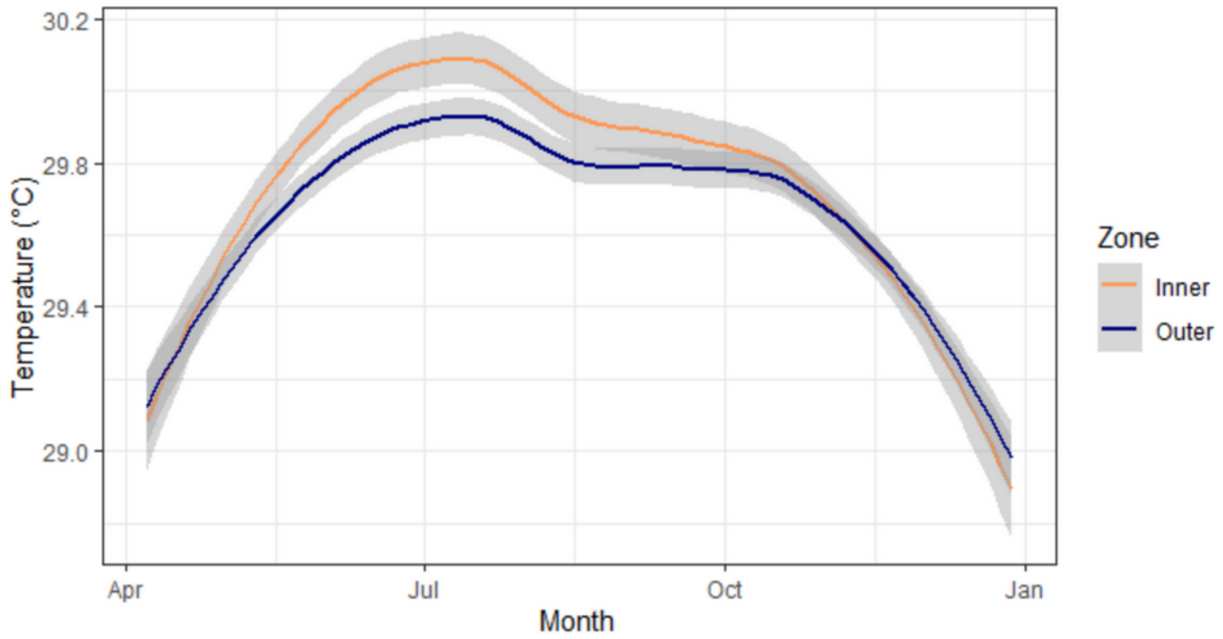


**Supplemental Figure 1.** Daily averages of seawater temperatures in the inner (orange) and outer (blue) zones of West Hagåtña Bay. The “geom_smooth” function in the R package ggplot2 was used to average the temperatures.

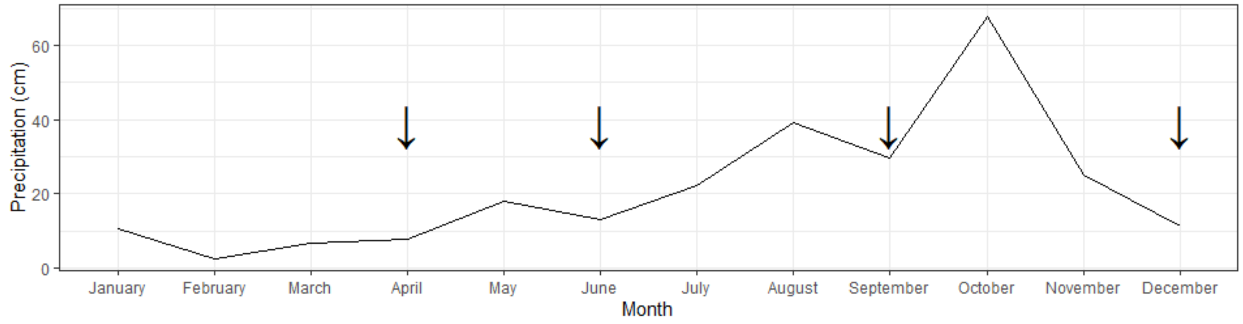


**Supplemental Figure 2.** Monthly rainfall throughout 2021. Arrows indicate sampling periods: the end of April, the end of June, the end of September, and the end of December.

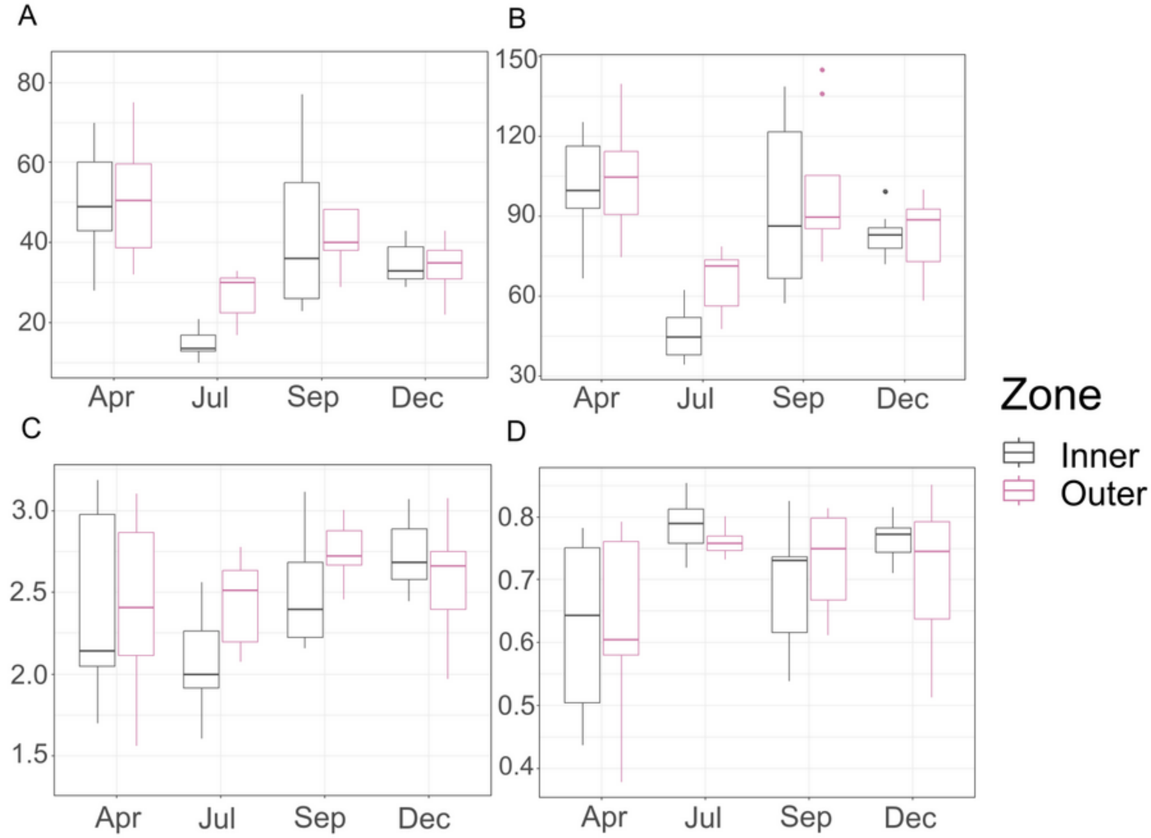


**Supplemental Figure 3.** Diversity measures of coral tissue. (A) Observed diversity; (B) phylogenetic diversity; (C) Shannon diversity; (D) evenness.

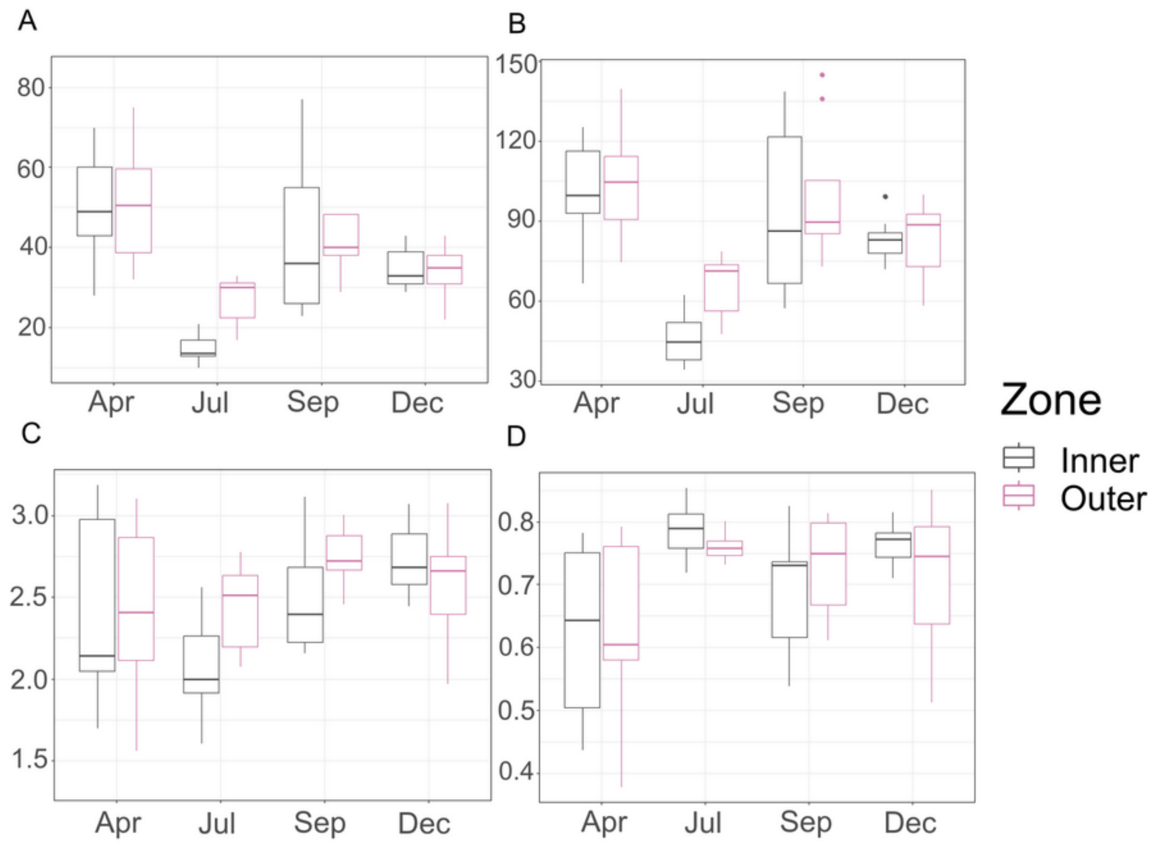


**Supplemental Figure 4.** Diversity measures of coral mucus. (A) Observed diversity; (B) phylogenetic diversity; (C) Shannon diversity; (D) evenness.

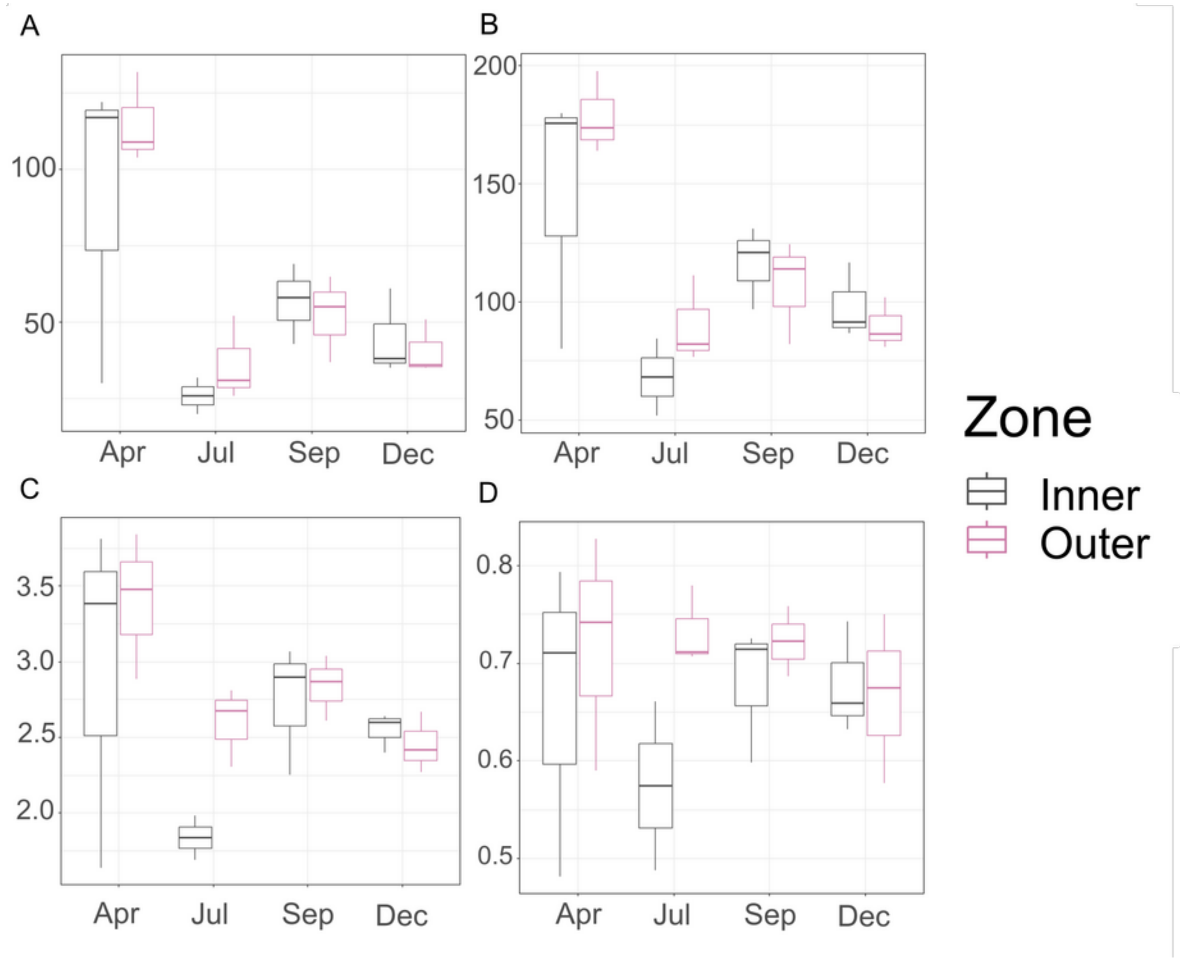


**Supplemental Figure 5.** Diversity measures of seawater. (A) Observed diversity; (B) phylogenetic diversity; (C) Shannon diversity; (D) evenness.
