## Supplementary material for "Seasonal dynamics and environmental drivers of tissue and mucus microbiomes in the staghorn coral *Acropora pulchra*": Data analysis scripts: FIB_analyses.R

```
library(readxl)
library(dplyr)
Water<-Water_Quality_Bacteriological_and_Chemical_Data_WAG <- read_excel("~/Graduate Research/Environmental Data/Water-Quality-Bacteriological-and-Chemical-Data-WAG.xlsx")
library(scales)
library(lubridate)
library(ggplot2)
library(gridExtra)

#Stats
Water$month<- month(ymd(Water$Date))

Water2 <- Water %>%
  group_by(month,In.Out)%>%
  summarise(Total.ColiformMean = mean(Total.Coliform),
            Total.ColiformMax = max(Total.Coliform),
            Total.ColiformMin = min(Total.Coliform),
            Total.ColiformRange = Total.ColiformMax - Total.ColiformMin,
            E.coliMean = mean(E.coli),
            E.coliMax = max(E.coli),
            E.coliMin = min(E.coli),
            E.coliRange = E.coliMax - E.coliMin,
            EnterococcusMean = mean(Enterococcus),
            EnterococcusMax = max(Enterococcus),
            EnterococcusMin = min(Enterococcus),
            EnterococcusRange = EnterococcusMax - EnterococcusMin)

# test normality with Shapiro-Wilk
shapiro.test(Water2$Total.ColiformMean) #normal

shapiro.test(Water2$E.coliMean) #normal

shapiro.test(Water2$EnterococcusMean) #normal

summary(aov(Total.ColiformMean ~ In.Out * month, data = Water2))

summary(aov(E.coliMean ~ In.Out * month, data = Water2))

summary(aov(EnterococcusMean ~ In.Out * month, data = Water2))

#Plots
ceiling_date(Water$Date,unit="month")
new_date <- as.Date(Water$Date, format = "%Y-/%m/-%d")
class(new_date)
month<-format(new_date,"%b")
Water$month <- month
class(Water$month)
sort(Water$month)
Water$month <- factor(Water$month, levels = month.abb)

#Enterococcus Plot
ep<-ggplot(Water,aes(x = month, y = Enterococcus,fill=In.Out))+
  geom_boxplot() +
  scale_fill_manual(values=c("red","lightblue"),name="Zone",labels=c("Inner", "Outer"))+
  xlab("Month")+
  ylab("log MPN/100mL")+
  ggtitle("Enterococcus Concentration")
ep
ep+ scale_y_continuous(trans = log10_trans()) +theme_bw()

#Coliform Plot
cp<-ggplot(Water,aes(x = month, y = Total.Coliform,fill=In.Out))+
  geom_boxplot() +
  scale_fill_manual(values=c("red", "lightblue"),name="Zone",labels=c("Inner", "Outer"))+
  xlab("Month")+
  ylab("log MPN/100mL")+
  ggtitle("Total Coliform Concentration")
cp
cp + scale_y_continuous(trans = log10_trans())+theme_bw()
                                                                            
#E. coli Plot
ecp<-ggplot(Water,aes(x = month, y = E.coli,fill=In.Out))+
  geom_boxplot() +
  scale_fill_manual(values=c("red", "lightblue"),name="Zone",labels=c("Inner", "Outer"))+
  xlab("Month")+
  ylab("log MPN/100mL")+
  ggtitle("E. coli Concentration")
ecp
ecp + scale_y_continuous(trans = log2_trans())+theme_bw()

grid.arrange(ep+ scale_y_continuous(trans = log10_trans()) +theme_bw(),cp + scale_y_continuous(trans = log10_trans())+theme_bw(),
             ecp + scale_y_continuous(trans = log10_trans())+theme_bw())
```
