## Supplementary material for "Seasonal dynamics and environmental drivers of tissue and mucus microbiomes in the staghorn coral *Acropora pulchra*": Data analysis scripts: Rain_plot.R

```
library(dplyr)
library(lubridate)
library(ggplot2)

rain<-read.csv("/Users/EPSCoR/Documents/Graduate Research/Environmental Data/Monthly_Rain_Data_2021.csv")

rain$month = factor(rain$Month, levels = month.name)
class(rain$Month)

rain$Month=1:12
names(rain$Month)=month.name

head(rain)

ggplot(rain,aes(x = month, y = Rain.cm.,group=1))+
  geom_line() +
  annotate("text", x = 4, y = 40, label = "↓", cex=10)+
  annotate("text", x = 6, y = 40, label = "↓", cex=10)+
  annotate("text", x = 9, y = 40, label = "↓", cex=10)+
  annotate("text", x = 12, y = 40, label = "↓", cex=10)+
  xlab("Month")+
  ylab("Precipitation (cm)")+
  theme_bw()

#Stats
daily<-read_excel("/Users/EPSCoR/Documents/Graduate Research/Environmental Data/Daily_Rain_Data.xlsx")

# test normality with Shapiro-Wilk
shapiro.test(daily$Precipitation) #not normal

kruskal.test(Precipitation ~ Month, data = daily)
```
