## Supplementary material for "Seasonal dynamics and environmental drivers of tissue and mucus microbiomes in the staghorn coral *Acropora pulchra*": Data analysis scripts: Temperature_analyses_plot.R

```
library(readxl)
library(tidyverse)
library(scales)
library(lubridate)
library(ggplot2)
Temperature <- read_excel("~/Graduate Research/Environmental Data/Temperature.xlsx", 
                          sheet = "AllTemps")
Temperature$month<- month(ymd(Temperature$Date))
class(month)

Temp2 <- Temperature %>%
  group_by(In.Out,Date,month)%>%
  summarise(TempMean = mean(Temp),
            TempMax = max(Temp),
            TempMin = min(Temp),
            TempRange = TempMax - TempMin) 
head(Temp2)
tail(Temp2)

# test normality with Shapiro-Wilk
shapiro.test(Temp2$TempMean) #not normal

kruskal.test(TempMean ~ In.Out, data = Temp2)

kruskal.test(TempMean ~ month, data = Temp2)

ggplot(Temp2,aes(x=Date,y=TempMean,color=In.Out))+
  geom_smooth()+
  xlab("Month")+
  ylab("Temperature (°C)")+
  scale_color_manual(values = c("#ff9955ff", "#00077eff"), name="Zone",labels=c("Inner", "Outer"))+
  theme_bw()
```
