## Supplementary material for "Seasonal dynamics and environmental drivers of tissue and mucus microbiomes in the staghorn coral *Acropora pulchra*": Data analysis scripts: BigData_merge.R

```
library("ggsci")
library('BiocManager')
library('Rcpp')
library(dada2)
library(phyloseq)
library(tidyverse)
library(dplyr)
library(usethis)
library(devtools)
#####################################

#####################################
library(ShortRead); packageVersion("ShortRead")
library(ggplot2); packageVersion("ggplot2")


# Merge multiple runs (if necessary)
st1 <- readRDS("/Users/EPSCoR/Documents/Graduate Research/Data Analysis/Apul-16s-run1/DADA2_Run1.rds")
st2 <- readRDS("/Users/EPSCoR/Documents/Graduate Research/Data Analysis/Apul-16s-run2/DADA2_Run2.rds")
st3 <- readRDS("/Users/EPSCoR/Documents/Graduate Research/Data Analysis/Apul-16s-run3/DADA2_Run3.rds")
st.all <- mergeSequenceTables(st1, st2, st3)
# Remove chimeras
seqtab <- removeBimeraDenovo(st.all, method="consensus", multithread=TRUE)
# Assign taxonomy
tax <- assignTaxonomy(seqtab, "/Users/EPSCoR/Documents/Graduate Research/Data Analysis/silva_nr99_v138_train_set.fa.gz", multithread=TRUE)
# Write to disk
saveRDS(seqtab, "/Users/EPSCoR/Documents/Graduate Research/Data Analysis/seqtab_final.rds") 
saveRDS(tax, "/Users/EPSCoR/Documents/Graduate Research/Data Analysis/tax_final.rds") 

######################################End BigData############################

#Let's inspect the taxonomic assignments:
taxa.print <- tax 
rownames(taxa.print) <- NULL
head(taxa.print)

write.csv(cbind(t(seqtab), tax), file ="/Users/EPSCoR/Documents/Graduate Research/Data Analysis/Apul-16S-BigData-taxa.csv")

################################
##### output ASV table #######
################################

ids <- paste0("ASV", seq(1, length(colnames(seqtab))))
colnames(seqtab)<-ids

write.csv(seqtab,file ="/Users/EPSCoR/Documents/Graduate Research/Data Analysis/OutputDADA_16S-v4_BigData.csv",quote=F)
write.csv(t(seqtab),file ="/Users/EPSCoR/Documents/Graduate Research/Data Analysis/tOutputDADA_16S-v4_BigData.csv",quote=F)

str(seqtab)

#Now, save outputs so can come back to the analysis stage at a later point if desired
saveRDS(seqtab, file="seqtab_nochim_16S-V4_BigData.rds")
saveRDS(tax, file="taxa_16S-V4_BigData.rds")

#If you need to read in previously saved data files
seqtab.nochim <- readRDS("/Users/EPSCoR/Documents/seqtab_nochim_16S-V4_BigData.rds")
taxa <- readRDS("/Users/EPSCoR/Documents/taxa_16S-V4_BigData.rds")

rownames(taxa.print) <- NULL
head(taxa.print)

# remove chloroplast
is.chloroplast <- tax[,"Order"] %in% "Chloroplast"
seqtable.nochloro <- seqtab.nochim[,!is.chloroplast]
dim(seqtable.nochloro)
taxonomy.nochloro <- tax[!is.chloroplast,]
dim(taxonomy.nochloro)

is.mitochondria <- taxonomy.nochloro[,"Family"] %in% "Mitochondria"
seqtable.nomito <- seqtable.nochloro[,!is.mitochondria]
taxonomy.nomito <- taxonomy.nochloro[!is.mitochondria,]
dim(seqtable.nomito)
dim(seqtable.nochloro)
dim(seqtab.nochim)

is.euk <- taxonomy.nomito[,"Kingdom"] %in% "Eukaryota" 
seqtable.noeuk <- seqtable.nomito[,!is.euk]
taxonomy.noeuk <- taxonomy.nomito[!is.euk,]
dim(seqtable.noeuk)
dim(seqtable.nomito)
dim(seqtable.nochloro)
dim(seqtab.nochim)

is.arch=taxonomy.noeuk[,"Kingdom"] %in% "Archaea"
seqtable.noarch <- seqtable.noeuk[,!is.arch]
taxonomy.noarch <- taxonomy.noeuk[!is.arch,]
dim(seqtable.noarch)
dim(seqtable.noeuk)
dim(seqtable.nomito)
dim(seqtable.nochloro)
dim(seqtab.nochim)
seqtab_nochim2 <- seqtable.noarch
tseqtab_nochim<-t(seqtab_nochim2)
dim(tseqtab_nochim)
taxonomy2 <- taxonomy.noarch
dim(taxonomy2)

ids <- paste0("ASV", seq(1, length(colnames(seqtab_nochim2))))
colnames(seqtab_nochim2)<-ids

ids <- paste0("ASV", seq(1, length(rownames(taxonomy2))))
rownames(taxonomy2)<-ids

saveRDS(t(seqtab_nochim2), file="/Users/EPSCoR/Documents/Graduate Research/Data Analysis/seqtab-min_BigData.rds")
saveRDS(taxonomy2, file="/Users/EPSCoR/Documents/Graduate Research/Data Analysis/tax-min_BigData.rds")

write.csv(seqtab_nochim2,file ="/Users/EPSCoR/Documents/Graduate Research/Data Analysis/seqtab-min_BigData.csv",quote=F)
write.csv(t(seqtab_nochim2),file ="/Users/EPSCoR/Documents/Graduate Research/Data Analysis/tseqtab-min_BigData.csv",quote=F)
write.csv(taxonomy2,file ="/Users/EPSCoR/Documents/Graduate Research/Data Analysis/taxo-min_BigData.csv",quote=F)


######################## PHYLOSEQ
library(phyloseq)
library(tidyverse)
library("dplyr")        # filter and reformat data frames

otu_mat<- read.table("/Users/EPSCoR/Documents/Graduate Research/Data Analysis/tseqtab-min_BigData.csv", sep=',', header=T,check.names = FALSE)
#Use check.names=FALSE to keep the numbered headings. Otherwise, R will relabel them with an "X" at the beginning
colnames(otu_mat)[1]<-"ASV"
colnames(otu_mat)[which(names(otu_mat)=="15T")]<-"15"


tax_mat<- read.table("/Users/EPSCoR/Documents/Graduate Research/Data Analysis/taxo-min_BigData.csv", sep=',', header=T)
colnames(tax_mat)[1]<-"ASV"

samples_df <- read.csv("/Users/EPSCoR/Documents/Graduate Research/Data Analysis/16S-Metadata-numberednames.csv", header=T)

row.names(otu_mat) <- otu_mat$ASV
colnames(otu_mat)[1]<-"ASV"

otu_mat <- otu_mat %>% select (-ASV) 

row.names(tax_mat) <- tax_mat$ASV
tax_mat <- tax_mat %>% select (-ASV)

row.names(samples_df) <- samples_df$SequenceID
samples_df <- samples_df %>% select (-SequenceID) 

otu_mat <- as.matrix(otu_mat)
tax_mat <- as.matrix(tax_mat)

OTU = otu_table(otu_mat, taxa_are_rows = TRUE)
TAX = tax_table(tax_mat)
samples = sample_data(samples_df)

carbom <- phyloseq(OTU, TAX, samples)
carbom
sample_names(carbom)
rank_names(carbom)
sample_variables(carbom)

total = median(sample_sums(carbom))
standf = function(x, t=total) round(t * (x / sum(x)))
carbom = transform_sample_counts(carbom, standf)

save.image("C:/Users/EPSCoR/Documents/Graduate Research/Data Analysis/BigData/BigData_merge.R.RData")


library("ape")
random_tree = rtree(ntaxa(carbom), rooted=TRUE, tip.label=taxa_names(carbom))
#plot(random_tree)
carbom = merge_phyloseq(carbom, samples, random_tree)

seqtab.all=data.frame(otu_table(carbom),check.names = FALSE)
write.csv(seqtab.all, file= "seqtab.all.csv")
seqtab.all=read.csv("seqtab.all.csv",row.names = 1)
library(MCMC.OTU)
seqtab.all.4mcmc=data.frame(t(otu_table(carbom)))


#Trim outliers#
seqtab.all.4mcmc=add_column(seqtab.all.4mcmc, sample = row.names(seqtab.all.4mcmc), .after = 0)
seqtab.all.trim=purgeOutliers(seqtab.all.4mcmc,count.columns=2:15524,sampleZcut=-2.5,otu.cut=0.0001,zero.cut=0.02)

write.csv(seqtab.all.trim, file= "seqtab.all.trim.csv")
seqtab.trim.4mcmc=t(seqtab.all.trim)

seqtab.trim.purge=seqtab.all.trim

seqtab.trim.purge.4mcmc<-t(seqtab.trim.purge)

ps.trim.purge<-phyloseq(otu_table(data.matrix(seqtab.trim.purge), taxa_are_rows=F), 
                        sample_data(samples_df), 
                        tax_table(tax_mat))
tissueps.trim.purge=subset_samples(ps.trim.purge, tissue.mucus.seawater=="tissue")
mucusps.trim.purge=subset_samples(ps.trim.purge, tissue.mucus.seawater=="mucus")
seawaterps.trim.purge=subset_samples(ps.trim.purge, tissue.mucus.seawater=="seawater")

write.csv(seqtab.trim.purge, file="/Users/EPSCoR/Documents/Graduate Research/Data Analysis/BigData/seqtab.trim.purge.csv")

#Get relative proportions of each taxon#
glom <- tax_glom(ps.trim.purge, taxrank = 'Phylum')
tax = tax_table(glom)[,"Phylum"]
otu = t(otu_table(glom))

table = merge(otu, tax,by=0,all=TRUE)
head(table)
rownames(table) = table[,"Phylum"]
table$Row.names = NULL
table$phylum=NULL
#write.csv(table, file ="/Users/EPSCoR/Documents/Graduate Research/Data Analysis/relabun_table_phylum.csv")

##For family##
glom2 <- tax_glom(ps.trim.purge, taxrank = 'Family')
tax2 = tax_table(glom2)[,"Family"]
otu2 = t(otu_table(glom2))

table2 = merge(otu2, tax2,by=0,all=TRUE)
head(table2)
rownames(table2) = table2[,"Family"]
table2$Row.names = NULL
table2$family=NULL
#write.csv(table2, file ="/Users/EPSCoR/Documents/Graduate Research/Data Analysis/relabun_table_family.csv")

#Separate by compartment
#Tissue
glom3 <- tax_glom(tissueps.trim.purge, taxrank = 'Phylum')
tax3 = tax_table(glom3)[,"Phylum"]
otu3 = t(otu_table(glom3))

table3 = merge(otu3, tax3,by=0,all=TRUE)
head(table3)
rownames(table3) = table3[,"Phylum"]
table3$Row.names = NULL
table3$phylum=NULL
write.csv(table3, file ="/Users/EPSCoR/Documents/Graduate Research/Data Analysis/tissue_relabun_table_phylum.csv")

#For family
glom4 <- tax_glom(tissueps.trim.purge, taxrank = 'Family')
tax4 = tax_table(glom4)[,"Family"]
otu4 = t(otu_table(glom4))

table4 = merge(otu4, tax4,by=0,all=TRUE)
head(table4)
rownames(table4) = table4[,"Family"]
table4$Row.names = NULL
table4$family=NULL
#write.csv(table4, file ="/Users/EPSCoR/Documents/Graduate Research/Data Analysis/tissue_relabun_table_family.csv")

#Mucus
glom5 <- tax_glom(mucusps.trim.purge, taxrank = 'Phylum')
tax5 = tax_table(glom5)[,"Phylum"]
otu5 = t(otu_table(glom5))

head(tax5)
head(otu5)

table5 = merge(otu5, tax5,by=0,all=TRUE)
head(table5)
rownames(table5) = table5[,"Phylum"]
table5$Row.names = NULL
table5$phylum=NULL
#write.csv(table5, file ="/Users/EPSCoR/Documents/Graduate Research/Data Analysis/mucus_relabun_table_phylum.csv")

#For family
glom6 <- tax_glom(mucusps.trim.purge, taxrank = 'Family')
tax6 = tax_table(glom6)[,"Family"]
otu6 = t(otu_table(glom6))

table6 = merge(otu6, tax6,by=0,all=TRUE)
head(table6)
rownames(table6) = table6[,"Family"]
table6$Row.names = NULL
table6$family=NULL
#write.csv(table6, file ="/Users/EPSCoR/Documents/Graduate Research/Data Analysis/mucus_relabun_table_family.csv")

#Seawater
glom7 <- tax_glom(seawaterps.trim.purge, taxrank = 'Phylum')
tax7 = tax_table(glom7)[,"Phylum"]
otu7 = t(otu_table(glom7))

head(tax7)
head(otu7)

table7 = merge(otu7, tax7,by=0,all=TRUE)
head(table7)
rownames(table7) = table7[,"Phylum"]
table7$Row.names = NULL
table7$phylum=NULL
#write.csv(table7, file ="/Users/EPSCoR/Documents/Graduate Research/Data Analysis/seawater_relabun_table_phylum.csv")

#For family
glom8 <- tax_glom(seawaterps.trim.purge, taxrank = 'Family')
tax8 = tax_table(glom8)[,"Family"]
otu8 = t(otu_table(glom8))

table8 = merge(otu8, tax8,by=0,all=TRUE)
head(table8)
rownames(table8) = table8[,"Family"]
table8$Row.names = NULL
table8$family=NULL
#write.csv(table8, file ="/Users/EPSCoR/Documents/Graduate Research/Data Analysis/seawater_relabun_table_family.csv")

samples_df <- read.csv("/Users/EPSCoR/Documents/Graduate Research/Data Analysis/16S-Metadata-numberednames.csv", header=T)

samples_thousand=samples_df

samples_thousand=add_column(samples_thousand, month = month.abb[samples_thousand$Month], .after = 0)
samples_thousand$month = factor(samples_thousand$month, levels = month.abb)
sort(samples_thousand$month)

#write.csv(samples_thousand, file="/Users/EPSCoR/Documents/Graduate Research/Data Analysis/samples.thousand.csv")
save.image("Processed_data.RData")
```
