## Supplementary material for "Seasonal dynamics and environmental drivers of tissue and mucus microbiomes in the staghorn coral *Acropora pulchra*": Data analysis scripts: DADA2_Run1.R

```
if (!requireNamespace("BiocManager", quietly = TRUE))
  install.packages("BiocManager")
#####################################

# installing packages: execute just once when first using this script:

library('BiocManager')
library('Rcpp')
library(dada2)
library(phyloseq)
packageVersion('phyloseq')

library(usethis)
library(devtools)
#####################################

#####################################
library(ShortRead); packageVersion("ShortRead")
library(ggplot2); packageVersion("ggplot2")

path <- "/Users/EPSCoR/Documents/Graduate Research/Data Analysis/Apul-16s-run1/"


list.files(path)

# Forward and reverse fastq filenames have format: SAMPLENAME_R1_001.fastq and SAMPLENAME_R2_001.fastq
fnFs <- sort(list.files(path, pattern="_R1_001.fastq", full.names = TRUE))
fnRs <- sort(list.files(path, pattern="_R2_001.fastq", full.names = TRUE))

# Extract sample names, assuming filenames have format: SAMPLENAME_XXX.fastq
sample.names <- sapply(strsplit(basename(fnFs), "_"), `[`, 1)

################## ################## ################## ################## 
################## 1. Inspect read quality profiles

plotQualityProfile(fnFs[1:2]) # check forward quality
plotQualityProfile(fnRs[1:2]) # check reverse quality

#Recommend trimming where quality profile crashes - in this case, >200 bases it gets below 30; still leaves ~50bp overlap
#If using this workflow on your own data: Your reads must still overlap after truncation in order to merge them later! If you are using a less-overlapping primer set, your truncLen must be large enough to maintain 20 + biological.length.variation nucleotides of overlap between them.
#BUT: For common ITS amplicon strategies, it is undesirable to truncate reads to a fixed length due to the large amount of length variation at that locus. That is OK, just leave out truncLen. Make sure you removed the forward and reverse primers from both the forward and reverse reads though! 

################## ################## ################## ################## 
################## 2. Filter and trim

# Place filtered files in filtered/ subdirectory

# Make directory and filenames for the filtered fastqs
filt_path <- file.path(path, "filtered")
if(!file_test("-d", filt_path)) dir.create(filt_path)
filtFs <- file.path(path, "filtered", paste0(sample.names, "_F_filt.fastq.gz"))
filtRs <- file.path(path, "filtered", paste0(sample.names, "_R_filt.fastq.gz"))

# Primers 16S region v4: Hyb515F and Hyb806R
FWD <- "GTGYCAGCMGCCGCGGTAA"  
REV <- "GGACTACHVGGGTWTCTAAT"

# Filter => Note, for OSU data, must trim b/c quality REALLY tanks later in the read; GSAF data fairly robust
names(filtFs) <- sample.names
names(filtRs) <- sample.names


out <- filterAndTrim(fnFs, filtFs, fnRs, filtRs, truncLen=c(185,185),
                     maxN=0, #DADA does not allow Ns
                     maxEE=c(2,2), #allow 1 expected errors, where EE = sum(10^(-Q/10)); more conservative, model converges
                     truncQ=2, 
                     trimLeft=c(19,20), #N nucleotides to remove from the start of each read: 16S-v4 primers: Hyb515F= 19bp; Hyb806R 20bp 
                     rm.phix=TRUE, #remove reads matching phiX genome
                     compress=TRUE, multithread=10,
                     matchIDs=TRUE,
                     ) # On Windows set multithread=FALSE
head(out)
tail(out)

################## ################## ################## ################## 
################## 3. learn the error rate 

setDadaOpt(MAX_CONSIST=30) #increase number of cycles to allow convergence # optional
errF <- learnErrors(filtFs, multithread=TRUE)
errR <- learnErrors(filtRs, multithread=TRUE)
plotErrors(errF, nominalQ=TRUE) #some issues with C2G and G2C variants being underestimated, but not terrible
plotErrors(errR, nominalQ=TRUE) #again, worse with G2C and C2G; a little T2G, but rest err on the side of being conservative (above red line)

################## ################## ################## ################## 
################## 4. Dereplicating #######

derepFs <- derepFastq(filtFs, verbose=TRUE)
derepRs <- derepFastq(filtRs, verbose=TRUE)
# Name the derep-class objects by the sample names
names(derepFs) <- sample.names
names(derepRs) <- sample.names

################## ################## ################## ################## 
################## 5. SAMPLE INFERENCE

dadaFs <- dada(filtFs, err=errF, multithread=TRUE)
dadaRs <- dada(filtRs, err=errR, multithread=TRUE)
dadaFs[[1]]

################## ################## ################## ################## 
################## 6. Merge paired reads

mergers <- mergePairs(dadaFs, filtFs, dadaRs, filtRs, verbose=TRUE)
# Inspect the merger data.frame from the first sample
head(mergers[[1]])
summary(mergers[[1]])

################## ################## ################## ################## 
################## 7. Construct sequence table

seqtab <- makeSequenceTable(mergers)
dim(seqtab)

# Inspect distribution of sequence lengths
table(nchar(getSequences(seqtab)))
plot(table(nchar(getSequences(seqtab)))) #real variants appear to be right in that "294-304" window


getN <- function(x) sum(getUniques(x))
track <- cbind(out, sapply(dadaFs, getN), sapply(dadaRs, getN), sapply(mergers, getN))
# If processing a single sample, remove the sapply calls: e.g. replace sapply(dadaFs, getN) with getN(dadaFs)
colnames(track) <- c("input", "filtered", "denoisedF", "denoisedR", "merged")
rownames(track) <- sample.names
head(track)

getN <- function(x) sum(getUniques(x))
track <- cbind(out, sapply(dadaFs, getN), sapply(dadaRs, getN), sapply(mergers, getN))
colnames(track) <- c("input", "filtered", "denoised", "merged", "tabled")
rownames(track) <- sample.names
head(track)
tail(track) 


saveRDS(seqtab, file="DADA2_run1.rds")

save.image("DADA2_Run1.RData")
```
