## Supplementary material for "Seasonal dynamics and environmental drivers of tissue and mucus microbiomes in the staghorn coral *Acropora pulchra*": Data analysis scripts: Diversity_analyses.R

```
load("~/Processed_Data.RData")

library(phyloseq)

row.names(samples_df)=samples_df$SequenceID
df.all <- data.frame(estimate_richness(ps.trim.purge, split=TRUE, measures=c("Shannon","InvSimpson","Observed")))

#Evenness#
H <- df.all$Shannon
S1 <- df.all$Observed
S <- log(S1)
evenness <- H/S
df.all$Evenness <- evenness
df.all

df.all$id <- rownames(df.all)
#seqtab.trim.purge<-seqtab.trim.purge %>% select(-cdat)


new.meta=samples_df[ (rownames(seqtab.trim.purge)), ]
library(tidyverse)

#write.csv(df.all,file="~/df.all.csv")
#write.csv(new.meta,file="~/new.meta.csv")

row.names(new.meta)
row.names(df.all)

df.all.div <- cbind(new.meta,df.all) 
df.all.div$id<-rownames(df.all.div)
row.names(df.all.div)

df.all$id <- rownames(df.all)
df.all.div <- cbind(df.all,new.meta) #add sample data

df.tissue <- data.frame(estimate_richness(tissueps.trim.purge, split=TRUE, measures=c("Shannon","InvSimpson","Observed")))
df.mucus <- data.frame(estimate_richness(mucusps.trim.purge, split=TRUE, measures=c("Shannon","InvSimpson","Observed")))
df.seawater <- data.frame(estimate_richness(seawaterps.trim.purge, split=TRUE, measures=c("Shannon","InvSimpson","Observed")))

df.tissue$id <- rownames(df.tissue)
Ht <- df.tissue$Shannon
S1t <- df.tissue$Observed
St <- log(S1t)
t.evenness <- Ht/St
df.tissue$Evenness <- t.evenness
df.tissue.div <- cbind(df.tissue,subset(new.meta, tissue.mucus.seawater=="tissue")) 

df.mucus$id <- rownames(df.mucus)
Hm <- df.mucus$Shannon
S1m <- df.mucus$Observed
Sm <- log(S1m)
m.evenness <- Hm/Sm
df.mucus$Evenness <- m.evenness
df.mucus.div <- cbind(df.mucus,subset(new.meta, tissue.mucus.seawater=="mucus")) 

df.seawater$id <- rownames(df.seawater)
Hs <- df.seawater$Shannon
S1s <- df.seawater$Observed
Ss <- log(S1s)
s.evenness <- Hs/Ss
df.seawater$Evenness <- s.evenness
df.seawater.div <- cbind(df.seawater,subset(new.meta, tissue.mucus.seawater=="seawater")) 

library(car)

#Faith's D
#install.packages("picante")
library("ape")
random_tree = rtree(ntaxa(ps.trim.purge), rooted=TRUE, tip.label=taxa_names(ps.trim.purge))
#plot(random_tree)
ps.trim.purge = merge_phyloseq(ps.trim.purge, samples, random_tree)

library("picante")

faithspd <- picante::pd(seqtab.trim.purge, random_tree)

ses.pd <- picante::ses.pd(seqtab.trim.purge, random_tree, 
                          null.model = c("taxa.labels", "richness", "frequency",
                          "sample.pool", "phylogeny.pool", "independentswap", "trialswap"),
                          runs = 999, iterations = 1000)
df.all.div <- cbind(df.all.div,faithspd,ses.pd)
pd.tissue <- subset(df.all.div, tissue.mucus.seawater=="tissue")
pd.mucus <-subset(df.all.div, tissue.mucus.seawater=="mucus")
pd.seawater <- subset(df.all.div, tissue.mucus.seawater=="seawater")

save.image("Diversity_analyses.RData")
```
