## Supplementary material for "Seasonal dynamics and environmental drivers of tissue and mucus microbiomes in the staghorn coral *Acropora pulchra*": Data analysis scripts: Diversity_Plots_final.R

```
load("C:/Users/EPSCoR/Documents/Diversity_analyses.RData")
library(phyloseq)
library(ggplot2); packageVersion("ggplot2")

#richness plots need to be done on phyloseq objects

#View(samples_df)
colnames(samples_df)[13]<-"Compartment"
colnames(samples_df)[3]<-"Zone"
samples_thousand <-within(samples_df, 
                    Compartment <- factor(Compartment,
                                                                levels = names(sort(table(Compartment),
                                                                                    decreasing=TRUE))))
df.all.div <-within(df.all.div, 
                          tissue.mucus.seawater <- factor(tissue.mucus.seawater,
                                                levels = names(sort(table(tissue.mucus.seawater),
                                                                    decreasing=TRUE))))

samples_thousand$Zone[samples_thousand$Zone == "In"] <- "Inner"
samples_thousand$Zone[samples_thousand$Zone == "Out"] <- "Outer"

ps.trim.purge<-phyloseq(otu_table(data.matrix(seqtab.trim.purge), taxa_are_rows=F), 
                        sample_data(samples_thousand), 
                        tax_table(tax_mat))
tissueps.trim.purge=subset_samples(ps.trim.purge, Compartment=="tissue")
mucusps.trim.purge=subset_samples(ps.trim.purge, Compartment=="mucus")
seawaterps.trim.purge=subset_samples(ps.trim.purge, Compartment=="seawater")

#Tissue
#PD
pd.tissue$Month <- factor(pd.tissue$Month, levels = c("Apr","Jul","Sep","Dec"))
ggplot(pd.tissue,aes(x = Month, y = PD, color = In.Out))+
  theme_bw()+
  theme(text = element_text(size = 20),  axis.text.x = element_text(angle = 0, vjust =.55 ))+
  scale_color_manual(name="Zone",values=c("#4d4d4dff","#CC79A7","#0055d4ff","#00077eff"),labels=c("Inner","Outer"))+xlab(NULL)+ylab(NULL)+ggtitle("Tissue PD by month and zone")+
  geom_boxplot(outlier.shape=NA)

#Shannon
ggplot(pd.tissue,aes(x = Month, y = Shannon, color = In.Out))+
  theme_bw()+
  theme(text = element_text(size = 20),  axis.text.x = element_text(angle = 0, vjust =.55 ))+
  scale_color_manual(name="Zone",values=c("#4d4d4dff","#CC79A7","#0055d4ff","#00077eff"),labels=c("Inner","Outer"))+xlab(NULL)+ylab(NULL)+ggtitle("Shannon diversity of tissue")+
  geom_boxplot(outlier.shape=NA)

#Observed
ggplot(pd.tissue,aes(x = Month, y = Observed, color = In.Out))+
  theme_bw()+
  theme(text = element_text(size = 20),  axis.text.x = element_text(angle = 0, vjust =.55 ))+
  scale_color_manual(name="Zone",values=c("#4d4d4dff","#CC79A7","#0055d4ff","#00077eff"),labels=c("Inner","Outer"))+xlab(NULL)+ylab(NULL)+ggtitle("Observed diversity of tissue")+
  geom_boxplot(outlier.shape=NA)

#Evenness
ggplot(pd.tissue,aes(x = Month, y = Evenness, color = In.Out))+
  theme_bw()+
  theme(text = element_text(size = 20),  axis.text.x = element_text(angle = 0, vjust =.55 ))+
  scale_color_manual(name="Zone",values=c("#4d4d4dff","#CC79A7","#0055d4ff","#00077eff"),labels=c("Inner","Outer"))+xlab(NULL)+ylab(NULL)+ggtitle("Evenness of tissue")+
  geom_boxplot(outlier.shape=NA)

#Mucus
#PD
ggplot(pd.mucus,aes(x = Month, y = PD, color = In.Out))+
  theme_bw()+
  theme(text = element_text(size = 20),  axis.text.x = element_text(angle = 0, vjust =.55 ))+
  scale_color_manual(name="Zone",values=c("#4d4d4dff","#CC79A7","#0055d4ff","#00077eff"),labels=c("Inner","Outer"))+xlab(NULL)+ylab(NULL)+ggtitle("Phylogenetic diversity of mucus")+
  geom_boxplot(outlier.shape=NA)

#Shannon
ggplot(pd.mucus,aes(x = Month, y = Shannon, color = In.Out))+
  theme_bw()+
  theme(text = element_text(size = 20),  axis.text.x = element_text(angle = 0, vjust =.55 ))+
  scale_color_manual(name="Zone",values=c("#4d4d4dff","#CC79A7","#0055d4ff","#00077eff"),labels=c("Inner","Outer"))+xlab(NULL)+ylab(NULL)+ggtitle("Shannon diversity of mucus")+
  geom_boxplot(outlier.shape=NA)

#Observed
ggplot(pd.mucus,aes(x = Month, y = Observed, color = In.Out))+
  theme_bw()+
  theme(text = element_text(size = 20),  axis.text.x = element_text(angle = 0, vjust =.55 ))+
  scale_color_manual(name="Zone",values=c("#4d4d4dff","#CC79A7","#0055d4ff","#00077eff"),labels=c("Inner","Outer"))+xlab(NULL)+ylab(NULL)+ggtitle("Observed diversity of mucus")+
  geom_boxplot(outlier.shape=NA)

#Evenness
ggplot(pd.mucus,aes(x = Month, y = Evenness, color = In.Out))+
  theme_bw()+
  theme(text = element_text(size = 20),  axis.text.x = element_text(angle = 0, vjust =.55 ))+
  scale_color_manual(name="Zone",values=c("#4d4d4dff","#CC79A7","#0055d4ff","#00077eff"),labels=c("Inner","Outer"))+xlab(NULL)+ylab(NULL)+ggtitle("Evenness of mucus")+
  geom_boxplot(outlier.shape=NA)


#Seawater
#PD
ggplot(pd.seawater,aes(x = Month, y = PD, color = In.Out))+
  theme_bw()+
  theme(text = element_text(size = 20),  axis.text.x = element_text(angle = 0, vjust =.55 ))+
  scale_color_manual(name="Zone",values=c("#4d4d4dff","#CC79A7","#0055d4ff","#00077eff"),labels=c("Inner","Outer"))+xlab(NULL)+ylab(NULL)+ggtitle("Seawater PD by month and zone")+
  geom_boxplot()

#Shannon
ggplot(pd.seawater,aes(x = Month, y = Shannon, color = In.Out))+
  theme_bw()+
  theme(text = element_text(size = 20),  axis.text.x = element_text(angle = 0, vjust =.55 ))+
  scale_color_manual(name="Zone",values=c("#4d4d4dff","#CC79A7","#0055d4ff","#00077eff"),labels=c("Inner","Outer"))+xlab(NULL)+ylab(NULL)+ggtitle("Shannon diversity of seawater")+
  geom_boxplot(outlier.shape=NA)

#Observed
ggplot(pd.seawater,aes(x = Month, y = Observed, color = In.Out))+
  theme_bw()+
  theme(text = element_text(size = 20),  axis.text.x = element_text(angle = 0, vjust =.55 ))+
  scale_color_manual(name="Zone",values=c("#4d4d4dff","#CC79A7","#0055d4ff","#00077eff"),labels=c("Inner","Outer"))+xlab(NULL)+ylab(NULL)+ggtitle("Observed diversity of seawater")+
  geom_boxplot(outlier.shape=NA)

#Evenness
ggplot(pd.seawater,aes(x = Month, y = Evenness, color = In.Out))+
  theme_bw()+
  theme(text = element_text(size = 20),  axis.text.x = element_text(angle = 0, vjust =.55 ))+
  scale_color_manual(name="Zone",values=c("#4d4d4dff","#CC79A7","#0055d4ff","#00077eff"),labels=c("Inner","Outer"))+xlab(NULL)+ylab(NULL)+ggtitle("Evenness of seawater")+
  geom_boxplot(outlier.shape=NA)

#All by compartment
plot_richness(ps.trim.purge, x="Compartment", measures=c("Shannon", "Observed"), color="Compartment") + theme_bw()+
  theme(text = element_text(size = 20), legend.position = "none", axis.text.x = element_text(angle = 0, vjust =.55 ))+
  scale_color_manual(name="Zone",values=c("#4d4d4dff","#0055d4ff","#87cddeff","#00077eff"))+xlab(NULL)+
  geom_boxplot()

ggplot(df.all.div, aes(x= tissue.mucus.seawater, y = PD, color = tissue.mucus.seawater))+
  theme(text = element_text(size = 20), legend.position = "none", axis.text.x = element_text(angle = 0, vjust =.55 ))+
  scale_color_manual(values=c("#4d4d4dff","#0055d4ff","#87cddeff","#00077eff"))+xlab(NULL)+
  geom_boxplot()+theme_bw()

ggplot(df.all.div, aes(x= tissue.mucus.seawater, y = Evenness, color = tissue.mucus.seawater))+
  theme(text = element_text(size = 20), legend.position = "none", axis.text.x = element_text(angle = 0, vjust =.55 ))+
  scale_color_manual(values=c("#4d4d4dff","#0055d4ff","#87cddeff","#00077eff"))+xlab(NULL)+
  geom_boxplot()+theme_bw()

ggplot(df.all.div, aes(x= tissue.mucus.seawater, y = Observed, color = tissue.mucus.seawater))+
  theme(text = element_text(size = 20), legend.position = "none", axis.text.x = element_text(angle = 0, vjust =.55 ))+
  scale_color_manual(values=c("#4d4d4dff","#0055d4ff","#87cddeff","#00077eff"))+xlab(NULL)+
  geom_boxplot()+theme_bw()

ggplot(df.all.div, aes(x= tissue.mucus.seawater, y = Shannon, color = tissue.mucus.seawater))+
  theme(text = element_text(size = 20), legend.position = "none", axis.text.x = element_text(angle = 0, vjust =.55 ))+
  scale_color_manual(values=c("#4d4d4dff","#0055d4ff","#87cddeff","#00077eff"))+xlab(NULL)+
  geom_boxplot()+theme_bw()

#Tissue by month
p<-plot_richness(tissueps.trim.purge, x="Month", measures=c("Shannon", "Observed"), color="Zone") + theme_bw()+
  theme(text = element_text(size = 20),  axis.text.x = element_text(angle = 0, vjust =.55 ))+
  scale_color_manual(name="Zone",values=c("#4d4d4dff","#CC79A7","#0055d4ff","#00077eff"),labels=c("Inner","Outer"))+xlab(NULL)+ylab(NULL)+ggtitle("Tissue by month and zone")+
  geom_boxplot(outlier.shape=NA)
p$layers[1]<-NULL
p

#Mucus by month
q<-plot_richness(mucusps.trim.purge, x="Month", measures=c("Shannon", "Observed"), color="Zone") + theme_bw()+
  theme(text = element_text(size = 20), axis.text.x = element_text(angle = 0, vjust =.55 ))+
  scale_color_manual(name="Zone",values=c("#4d4d4dff","#CC79A7","#0055d4ff","#00077eff"),labels=c("Inner","Outer"))+xlab(NULL)+ylab(NULL)+ggtitle("Mucus by month and zone")+
  geom_boxplot(outlier.shape=NA)
q$layers[1]<-NULL
q

#Seawater by month
r<-plot_richness(seawaterps.trim.purge, x="Month", measures=c("Shannon", "Observed"), color="Zone") + theme_bw()+
  theme(text = element_text(size = 20),  axis.text.x = element_text(angle = 0, vjust =.55 ))+
  scale_color_manual(name="Zone",values=c("#4d4d4dff","#CC79A7","#0055d4ff","#00077eff"),labels=c("Inner","Outer"))+xlab(NULL)+ylab(NULL)+ggtitle("Seawater by month and zone")+
  geom_boxplot()
r$layers[1]<-NULL
r
library(gridExtra)
grid.arrange(p,q,r)

#Unifrac
#Create a phylogenetic tree with the ape package

library("ape")
random_tree = rtree(ntaxa(ps.trim.purge), rooted=TRUE, tip.label=taxa_names(ps.trim.purge))
#plot(random_tree)
ps.trim.purge=merge_phyloseq(ps.trim.purge,random_tree)
ps.trim.purge

#Weighted Unifrac
carbom.ord <- ordinate(ps.trim.purge,"PCoA", "unifrac",weighted=T)

wt.unifrac.ssn <- plot_ordination(ps.trim.purge, carbom.ord, shape="Season", color = "Compartment")
wt.unifrac.ssn <- wt.unifrac.ssn + ggtitle("Weighted UniFrac") + geom_point(size = 3)
wt.unifrac.ssn <- wt.unifrac.ssn + theme_classic() 
print(wt.unifrac.ssn)
print(wt.unifrac.ssn + stat_ellipse(linewidth=1))

wt.unifrac.ste <- plot_ordination(ps.trim.purge, carbom.ord, shape="Zone", color = "Compartment")
wt.unifrac.ste <- wt.unifrac.ste + ggtitle("Weighted UniFrac") + geom_point(size = 3)
wt.unifrac.ste <- wt.unifrac.ste + theme_classic() 
print(wt.unifrac.ste)
print(wt.unifrac.ste + stat_ellipse(linewidth=1))
library(gridExtra)
grid.arrange(wt.unifrac.ssn+ stat_ellipse(linewidth=1), wt.unifrac.ste+ stat_ellipse(linewidth=1))
save.image("~/Diversity_Plots_final.RData")
```
