## Supplementary material for "Seasonal dynamics and environmental drivers of tissue and mucus microbiomes in the staghorn coral *Acropora pulchra*": Data analysis scripts: RelativeAbundancePlots_final.R

```
######################## PHYLOSEQ
library(ggplot2)
library(ggsci)
library(phyloseq)
library(tidyverse)

library("dplyr")    
load("~/Diversity_analyses.RData")

#Separate by family#
glom <- tax_glom(ps.trim.purge, taxrank = 'Family')
glom # should list taxa as families

data <- psmelt(glom) # create dataframe from phyloseq object
data$family <- as.character(data$Family) #convert to character

#Taxonomic filtering
table(tax_table(ps.trim.purge)[, "Family"], exclude = NULL)
#Shows number of features of each taxa
ps <- subset_taxa(ps.trim.purge, !is.na(Family) & !Family %in% c())
#Prevalence filtering
# Compute prevalence of each feature, store as data.frame
prevdf <- apply(X = otu_table(ps),
                MARGIN = ifelse(taxa_are_rows(ps), yes = 1, no = 2),
                FUN = function(x){sum(x > 0)})
# Add taxonomy and total read counts to this data.frame
prevdf <- data.frame(Prevalence = prevdf,
                     TotalAbundance = taxa_sums(ps),
                     tax_table(ps))
#Compute total and averages of each prevalence
plyr::ddply(prevdf, "Family", function(df1){cbind(mean(df1$Prevalence),sum(df1$Prevalence))})

# Filter entries with unidentified Family.
ps <- subset_taxa(ps, !Family %in% prevdf)
ps

# Subset to the remaining families
prevdf1 <- subset(prevdf, Family %in% get_taxa_unique(ps, "Family"))

# Define prevalence threshold as 5% of total samples
prevalenceThreshold <- 0.05 * nsamples(ps)

# Execute prevalence filter, using `prune_taxa()` function
keepTaxa <- rownames(prevdf1)[(prevdf1$Prevalence >= prevalenceThreshold)]
ps1 <- prune_taxa(keepTaxa, ps)
ps1

ps1_family <- tax_glom(ps1, "Family", NArm = TRUE)
# Get top 10 families
top10_family <- names(sort(taxa_sums(ps1_family), decreasing=TRUE))[1:10]

# Transform Taxa counts to relative abundance
ps1_family_relabun <- transform_sample_counts(ps1_family, function(ASV) ASV/sum(ASV) * 100)

# Extract the top 10 taxa
ps1_family_top10 <- prune_taxa(top10_family, ps1_family_relabun)

# Convert into dataframe
taxa_abundance_table_family <- psmelt(ps1_family_top10)

#Exclude by zone and compartment
taxa_abundance_table_family$In.Out[taxa_abundance_table_family$Zone == "In"] <- "Inner"
taxa_abundance_table_family$In.Out[taxa_abundance_table_family$Zone == "Out"] <- "Outer"
taxa_abundance_table_family$Month <- month.abb[taxa_abundance_table_family$Month]
taxa_abundance_table_family$Month = factor(taxa_abundance_table_family$Month, levels = month.abb)

seawater2<-filter(taxa_abundance_table_family,tissue.mucus.seawater=="seawater")
tissue2<-filter(taxa_abundance_table_family,tissue.mucus.seawater=="tissue")
mucus2<-filter(taxa_abundance_table_family,tissue.mucus.seawater=="mucus")

#Plot the family
Tissue_family <- tissue2 %>%
  ggplot(aes(x =Sample, y = Abundance, fill = Family)) +
  geom_bar(
    #position="fill", 
    stat = "identity") +
  labs(x = "Samples (n = 70)",
       y = "Relative Abundance",
       title = "Tissue Family Relative Abundance") +
  facet_grid(~Month + In.Out, scales = "free") +
  theme(
    axis.text.x = element_text(size = 10, angle = 90, vjust = 0.5, hjust = 1),
    axis.text.y = element_text(size = 12),
    legend.text = element_text(size = 10),
    strip.text = element_text(size = 12)
  )
Tissue_family+scale_fill_npg()

Mucus_family <- mucus2 %>%
  ggplot(aes(x =Sample, y = Abundance, fill = Family)) +
  geom_bar(
    stat = "identity") +
  labs(x = "Samples (n = 68)",
       y = "Relative Abundance",
       title = "Mucus Family Relative Abundance") +
  facet_grid(~Month + In.Out, scales = "free") +
  theme(
    axis.text.x = element_text(size = 10, angle = 90, vjust = 0.5, hjust = 1),
    axis.text.y = element_text(size = 12),
    legend.text = element_text(size = 10),
    strip.text = element_text(size = 12)
  )
Mucus_family+scale_fill_npg()

Seawater_family <- seawater2 %>%
  ggplot(aes(x =Sample, y = Abundance, fill = Family)) +
  geom_bar(
    #position="fill", 
    stat = "identity") +
  labs(x = "Samples (n = 23)",
       y = "Relative Abundance",
       title = "Seawater Family Relative Abundance") +
  facet_grid(~Month + In.Out, scales = "free") +
  theme(
    axis.text.x = element_text(size = 10, angle = 90, vjust = 0.5, hjust = 1),
    axis.text.y = element_text(size = 12),
    legend.text = element_text(size = 10),
    strip.text = element_text(size = 12),
  )
Seawater_family+scale_fill_npg()

library(gridExtra)
grid.arrange(Tissue_family+scale_fill_npg(),Mucus_family+scale_fill_npg(),Seawater_family+scale_fill_npg())
```
