## Supplementary material for "Seasonal dynamics and environmental drivers of tissue and mucus microbiomes in the staghorn coral *Acropora pulchra*": Data analysis scripts: Statistical_Analyses.R

```
#Statistical Testing#
load("~/Diversity_analyses.RData")
library(phyloseq)
library(ggplot2)
library(agricolae)

#Looking at Shannon Diversity
#Normality and Heteroskedasticity: 
###For post-hoc ANOVAs###
tissue.mucus <- subset(df.all.div,tissue.mucus.seawater=="tissue" | tissue.mucus.seawater=="mucus")
tissue.seawater <- subset(df.all.div,tissue.mucus.seawater=="tissue" | tissue.mucus.seawater=="seawater")
mucus.seawater <- subset(df.all.div,tissue.mucus.seawater=="seawater" | tissue.mucus.seawater=="mucus")

library(car)

#Testing in vs out
shapiro.test(df.all.div$Observed) 

shapiro.test(df.all.div$Shannon) 
#Data is not normally distributed
#Now Dunn Test
library(DescTools)
DescTools::DunnTest(Shannon ~ tissue.mucus.seawater, df.all.div)

hist(df.all.div$Shannon)

shapiro.test(df.all.div$Observed) #use Dunn Test
#W = 0.82598, p-value = 1.446e-12
DescTools::DunnTest(Observed ~ tissue.mucus.seawater, df.all.div)

shapiro.test(df.tissue.div$Shannon) #not normal

shapiro.test(df.mucus.div$Shannon) #not normal

shapiro.test(df.seawater.div$Shannon) #normal


shapiro.test(df.all.div$Evenness)
#not normal
DescTools::DunnTest(Evenness ~ tissue.mucus.seawater, df.all.div)

shapiro.test(df.all.div$PD)
#not normal
DescTools::DunnTest(PD ~ tissue.mucus.seawater, df.all.div)

#################################################################
###Look at In.Out and Month interactions##
#################################################################

summary(aov(Shannon ~ In.Out*Month, data = df.seawater.div))

#################################################################
###Kruskal-Wallis tests: overall##
#################################################################
#Make new column for interaction of compartment, zone, and month for post-hoc dunn test#
df.all.div$com.zone.month <-paste(df.all.div$tissue.mucus.seawater,df.all.div$In.Out,df.all.div$month)
df.all.div$com.zone <- paste(df.all.div$tissue.mucus.seawater,df.all.div$In.Out)
df.all.div$com.month<-paste(df.all.div$tissue.mucus.seawater,df.all.div$Month)
df.all.div$zone.month<-paste(df.all.div$Month,df.all.div$In.Out)

kruskal.test(Shannon ~ tissue.mucus.seawater, data=df.all.div)
#significant
DescTools::DunnTest(Shannon ~ tissue.mucus.seawater, df.all.div)

DescTools::DunnTest(Observed ~ tissue.mucus.seawater, df.all.div)

#Shannon by zone#
kruskal.test(Shannon ~ In.Out, data = df.all.div)
#not significant

#Shannon by Month#
kruskal.test(Shannon ~ Month, data = df.all.div)

#Shannon by zone and month#
kruskal.test(Shannon ~ zone.month, df.all.div)
#no

kruskal.test(Shannon ~ com.zone, df.all.div)

kruskal.test(Shannon ~ com.month, df.all.div)

kruskal.test(Shannon ~ com.zone.month, df.all.div)

#Evenness
shapiro.test(df.all.div$Evenness)#not normal
kruskal.test(Evenness ~ tissue.mucus.seawater, data=df.all.div)

kruskal.test(Evenness ~ In.Out, data = df.all.div)

kruskal.test(Evenness ~ Month, data = df.all.div)

kruskal.test(Evenness ~ zone.month, df.all.div)

kruskal.test(Evenness ~ com.zone.month, data = df.all.div)

DunnTest(Evenness ~ com.zone.month, df.all.div)

kruskal.test(Evenness ~ com.zone, df.all.div)

kruskal.test(Evenness ~ com.month, df.all.div)

#PD
shapiro.test(df.all.div$PD)#not normal
kruskal.test(PD ~ tissue.mucus.seawater, data=df.all.div)

kruskal.test(PD ~ In.Out, data = df.all.div)

kruskal.test(PD ~ Month, data = df.all.div)

kruskal.test(PD ~ zone.month, df.all.div)

kruskal.test(PD ~ com.zone.month, data = df.all.div)

DunnTest(PD ~ com.zone.month, df.all.div)

kruskal.test(PD ~ com.zone, df.all.div)

DunnTest(PD ~ com.zone, df.all.div)

kruskal.test(PD ~ com.month, df.all.div)

DunnTest(PD ~ com.month, df.all.div)

#Interaction of compartment:zone:month
kruskal.test(Shannon ~ com.zone.month, data = df.all.div)

DunnTest(Shannon ~ com.zone.month, df.all.div)

kruskal.test(Shannon ~ com.zone, df.all.div)

DunnTest(Shannon ~ com.zone, df.all.div)

kruskal.test(Shannon ~ com.month, df.all.div)


#################################################################
###Kruskal-Wallis tests: tissue##
#################################################################
#Make new column for interaction of compartment, zone, and month for post-hoc dunn test#
df.tissue.div$zone.month<-paste(df.tissue.div$Month,df.tissue.div$In.Out)
pd.tissue$zone.month<-paste(pd.tissue$Month,pd.tissue$In.Out)

kruskal.test(Shannon ~ In.Out, data = df.tissue.div)

kruskal.test(Shannon ~ Month, data = df.tissue.div)

kruskal.test(Shannon ~ zone.month, df.tissue.div)

shapiro.test(df.tissue.div$Evenness)#not normal

kruskal.test(Evenness ~ In.Out, data = df.tissue.div)

kruskal.test(Evenness ~ Month, data = df.tissue.div)

kruskal.test(Evenness ~ zone.month, df.tissue.div)

shapiro.test(pd.tissue$PD)#not normal
kruskal.test(PD ~ In.Out, data = pd.tissue)

kruskal.test(PD ~ Month, data = pd.tissue)

kruskal.test(PD ~ zone.month, pd.tissue)

#################################################################
###Kruskal-Wallis tests: mucus##
#################################################################
#Make new column for interaction of compartment, zone, and month for post-hoc dunn test#
df.mucus.div$zone.month<-paste(df.mucus.div$Month,df.mucus.div$In.Out)
pd.mucus$zone.month<-paste(pd.mucus$Month,pd.mucus$In.Out)

kruskal.test(Shannon ~ In.Out, data = df.mucus.div)

kruskal.test(Shannon ~ Month, data = df.mucus.div)

DunnTest(Shannon ~ Month, df.mucus.div)

kruskal.test(Shannon ~ zone.month, df.mucus.div)

DunnTest(Shannon ~ zone.month, df.mucus.div)

shapiro.test(df.mucus.div$Evenness)#not normal

kruskal.test(Evenness ~ In.Out, data = df.mucus.div)

kruskal.test(Evenness ~ Month, data = df.mucus.div)

DunnTest(Evenness ~ Month, df.mucus.div)

kruskal.test(Evenness ~ zone.month, df.mucus.div)

DunnTest(Evenness ~ zone.month, df.mucus.div)

shapiro.test(pd.mucus$PD)#normal

hist(pd.mucus$PD)

summary(aov(PD ~ In.Out, data = pd.mucus))

summary(aov(PD ~ Month, data = pd.mucus))

summary(aov(PD ~ zone.month, pd.mucus))

#################################################################
###Kruskal-Wallis tests: seawater##
#################################################################
#Make new column for interaction of compartment, zone, and month for post-hoc dunn test#
df.seawater.div$zone.month<-paste(df.seawater.div$Month,df.seawater.div$In.Out)
pd.seawater$zone.month<-paste(pd.seawater$Month,pd.seawater$In.Out)

shapiro.test(pd.seawater$PD)#not normal

kruskal.test(PD ~  In.Out, data = pd.seawater)

kruskal.test(PD ~ Month, data = pd.seawater)

kruskal.test(PD ~ zone.month, pd.seawater)

#Evenness#
shapiro.test(df.seawater.div$Evenness)#normal
summary(aov(Evenness ~In.Out, data = pd.seawater))

summary(aov(Evenness ~ Month, data = pd.seawater))

summary(aov(Evenness ~ zone.month, pd.seawater))

summary(aov(PD ~ In.Out, data = pd.seawater))


#Beta Diversity
load("~/Processed_Data.RData")

library("ape")
random_tree = rtree(ntaxa(ps.trim.purge), rooted=TRUE, tip.label=taxa_names(ps.trim.purge))
#plot(random_tree)
ps.all.trim_tree = merge_phyloseq(ps.trim.purge, samples, random_tree)
ps.all.trim_tree.ord <- ordinate(ps.all.trim_tree,"PCoA", "unifrac",weighted=T)

#Permanova
library(vegan)

metadf<-data.frame(sample_data(ps.all.trim_tree))
unifrac.dist<-UniFrac(ps.all.trim_tree,
                      weighted=T,
                      normalized=T,
                      parallel=F,
                      fast=T)
library(pairwiseAdonis)
perma<-adonis2(unifrac.dist~tissue.mucus.seawater*In.Out*Month, data=metadf)
perma 

pairwise.adonis(unifrac.dist,factors=metadf$tissue.mucus.seawater,perm=1000)

pairwise.adonis(unifrac.dist,factors=metadf$Month,perm=1000)

#Separate compartments
#Beta Diversity
random_tree = rtree(ntaxa(tissueps.trim.purge), rooted=TRUE, tip.label=taxa_names(tissueps.trim.purge))
#plot(random_tree)
tissue_tree = merge_phyloseq(tissueps.trim.purge, samples, random_tree)
tissue_tree.ord <- ordinate(tissue_tree,"PCoA", "unifrac",weighted=T)

#Permanova
metadf<-data.frame(sample_data(tissue_tree))
unifrac.dist<-UniFrac(tissue_tree,
                      weighted=T,
                      normalized=T,
                      parallel=F,
                      fast=T)
perma<-adonis2(unifrac.dist~In.Out*Month, data=metadf)
perma
pairwise.adonis(unifrac.dist,factors=metadf$Month,perm=1000)

#Mucus
random_tree = rtree(ntaxa(mucusps.trim.purge), rooted=TRUE, tip.label=taxa_names(mucusps.trim.purge))
#plot(random_tree)
mucus_tree = merge_phyloseq(mucusps.trim.purge, samples, random_tree)
mucus_tree.ord <- ordinate(mucus_tree,"PCoA", "unifrac",weighted=T)

#Permanova
metadf<-data.frame(sample_data(mucus_tree))
unifrac.dist<-UniFrac(mucus_tree,
                      weighted=T,
                      normalized=T,
                      parallel=F,
                      fast=T)
perma<-adonis2(unifrac.dist ~ Month*In.Out, data=metadf)
perma 

pairwise.adonis(unifrac.dist,factors=metadf$Month,perm=1000)

#Seawater
random_tree = rtree(ntaxa(seawaterps.trim.purge), rooted=TRUE, tip.label=taxa_names(seawaterps.trim.purge))
#plot(random_tree)
seawater_tree = merge_phyloseq(seawaterps.trim.purge, samples, random_tree)
seawater_tree.ord <- ordinate(seawater_tree,"PCoA", "unifrac",weighted=T)

#Permanova
metadf<-data.frame(sample_data(seawater_tree))
unifrac.dist<-UniFrac(seawater_tree,
                      weighted=T,
                      normalized=T,
                      parallel=F,
                      fast=T)
perma<-adonis2(unifrac.dist~In.Out*Month, data=metadf)
perma 

pairwise.adonis(unifrac.dist,factors=metadf$Month,perm=1000)
```
