## Supplementary material for "Seasonal dynamics and environmental drivers of tissue and mucus microbiomes in the staghorn coral *Acropora pulchra*": Data analysis scripts: README.docx

Scripts used for analysis of environmental data, microbiome diversity and composition, and the construction of the structural equation model (SEM).

The SEM relies on data and products found in the environmental data and microbiome data analysis folders.
