## Supplementary material for "Seasonal dynamics and environmental drivers of tissue and mucus microbiomes in the staghorn coral *Acropora pulchra*": Data analysis scripts: Enviro_Data_Combined_for_SEM.R

```
library(readxl)
library(dplyr)
library(lubridate)
library(readr)

Temperature <- read_excel("~/Graduate Research/Environmental Data/Temperature.xlsx", 
                          sheet = "AllTemps")
Temperature$month<- month(ymd(Temperature$Date))
Temp2 <- Temperature %>%
  group_by(month,In.Out)%>%
  summarise(TempMean = mean(Temp),
            TempMax = max(Temp),
            TempMin = min(Temp),
            TempRange = TempMax - TempMin)
head(Temp2)


Rain <- read_csv("~/Graduate Research/Environmental Data/Monthly_Rain_Data_2021.csv")
names(Rain)[3]<-"Rain"
Rain$month<-c(1,2,3,4,5,6,7,8,9,10,11,12)

Rain <- Rain %>% select(-Month, -Day)
head(Rain)

FIB <- read_excel("~/Graduate Research/Environmental Data/Water-Quality-Bacteriological-and-Chemical-Data-WAG.xlsx")
FIB$month<- month(ymd(FIB$Date))

FIB2 <- FIB %>%
  group_by(month,In.Out)%>%
  summarise(Total.ColiformMean = mean(Total.Coliform),
            Total.ColiformMax = max(Total.Coliform),
            Total.ColiformMin = min(Total.Coliform),
            Total.ColiformRange = Total.ColiformMax - Total.ColiformMin,
            E.coliMean = mean(E.coli),
            E.coliMax = max(E.coli),
            E.coliMin = min(E.coli),
            E.coliRange = E.coliMax - E.coliMin,
            EnterococcusMean = mean(Enterococcus),
            EnterococcusMax = max(Enterococcus),
            EnterococcusMin = min(Enterococcus),
            EnterococcusRange = EnterococcusMax - EnterococcusMin)

FIB2$In.Out[FIB2$In.Out == "IN"] <- "In"
FIB2$In.Out[FIB2$In.Out == "OUT"] <- "Out"

load("~/Diversity_analyses.RData")

names(df.all.div)[12] <- "month"

df.new <- df.all.div %>%
  group_by(month,tissue.mucus.seawater,In.Out,SequenceID)%>%
  summarise(ShanMean = mean(Shannon),
            ShanMax = max(Shannon),
            ShanMin = min(Shannon),
            ShanRange = ShanMax - ShanMin)

#To combine all of these environmental data frames, use full_join
FIB_Temp<-full_join(FIB2,Temp2, by = c("month","In.Out"))

df.FT<-full_join(FIB_Temp,df.new, by = c("month","In.Out"))

df.FTR<-full_join(df.FT,Rain, by = "month")

new_FTR<-subset(df.FTR,  month=="4" | month=="7"| month=="9"| month=="12")
View(new_FTR)

seawater<-filter(new_FTR, tissue.mucus.seawater=="seawater")
mucus<-filter(new_FTR, tissue.mucus.seawater=="mucus")
tissue<-filter(new_FTR, tissue.mucus.seawater=="tissue")

save.image("~/Enviro_Data_Combined_for_SEM.RData")
```
