## Supplementary material for "Seasonal dynamics and environmental drivers of tissue and mucus microbiomes in the staghorn coral *Acropora pulchra*": Data analysis scripts: SEM_analyses.R

```
load("~/Enviro_Data_Combined_for_SEM.RData")

#Remove month from all these analyses#

library(piecewiseSEM)
library(nlme)
library(ggplot2)
library(effects)
library(dplyr)

FTR_models_colinearity <- list(
  Coliform.mod2 = lme(Total.ColiformMean ~ Rain + TempMean, random = ~1|In.Out, data= new_FTR),
  E.coli.mod2 = lme(E.coliMean ~ Rain + TempMean, random = ~1|In.Out, data = new_FTR),
  Entero.mod2 = lme(EnterococcusMean ~ Rain + TempMean, random = ~1|In.Out, data = new_FTR),
  ###Correlated error structures
  Total.ColiformMean %~~% E.coliMean, Total.ColiformMean %~~% EnterococcusMean,
  E.coliMean %~~% Total.ColiformMean, E.coliMean %~~% EnterococcusMean,
  EnterococcusMean %~~% Total.ColiformMean, EnterococcusMean %~~% E.coliMean,
  Shan_index.mod = lm(ShanMean ~ Total.ColiformMean + E.coliMean + EnterococcusMean, data = new_FTR))
FTR.co.psem <- as.psem(FTR_models_colinearity)
summary(FTR.co.psem, .direction = T)

#Tissue
tissue.FTR_models_colinearity <- list(
  Coliform.mod2 = lme(Total.ColiformMean ~ Rain + TempMean, random = ~1|In.Out, data = tissue),
  E.coli.mod2 = lme(E.coliMean ~ Rain + TempMean, random = ~1|In.Out, data = tissue),
  Entero.mod2 = lme(EnterococcusMean ~ Rain + TempMean, random = ~1|In.Out, data = tissue),
  ###Correlated error structures
  Total.ColiformMean %~~% E.coliMean, Total.ColiformMean %~~% EnterococcusMean,
  E.coliMean %~~% Total.ColiformMean, E.coliMean %~~% EnterococcusMean,
  EnterococcusMean %~~% Total.ColiformMean, EnterococcusMean %~~% E.coliMean,
  Shan_index.mod = lm(ShanMean ~ Total.ColiformMean + E.coliMean + EnterococcusMean, data = tissue))
FTR.co.tissue <- as.psem(tissue.FTR_models_colinearity)
(FTR.co.tissue.summary <- summary(FTR.co.tissue, .progressBar = T))

#Mucus
mucus.FTR.co_models <- list(
  Coliform.mod2 = lme(Total.ColiformMean ~ Rain + TempMean, random = ~1|In.Out, data = mucus),
  E.coli.mod2 = lme(E.coliMean ~ Rain + TempMean, random = ~1|In.Out, data = mucus),
  Entero.mod2 = lme(EnterococcusMean ~ Rain + TempMean, random = ~1|In.Out, data = mucus),
  ###Correlated error structures
  Total.ColiformMean %~~% E.coliMean, Total.ColiformMean %~~% EnterococcusMean,
  E.coliMean %~~% Total.ColiformMean, E.coliMean %~~% EnterococcusMean,
  EnterococcusMean %~~% Total.ColiformMean, EnterococcusMean %~~% E.coliMean,
  ShanMean %~~% Rain,
  Shan_index.mod = lm(ShanMean ~ Total.ColiformMean + E.coliMean + EnterococcusMean, data = mucus))

mucus.co.psem <- as.psem(mucus.FTR.co_models)
summary(mucus.co.psem)
```
